# Metabolism fine tuning and cardiokines secretion represent adaptative responses of the heart to High Fat and High Sugar Diets in flies

**DOI:** 10.1101/2024.10.09.617032

**Authors:** Lucie Khamvongsa-Charbonnier, Laurent Kremmer, Magali Torres, Sallouha Krifa, Charis Aubert, Alice Corbet, Loïc Crespo, Laurence Röder, Laurent Perrin, Nathalie Arquier

## Abstract

Cardiopathies are one of the leading causes of death in obese diabetics. Diabetic cardiomyopathies are notably characterized by contractile dysfunctions. Using the *Drosophila* model for cardiac function in pathophysiological context, we identified a set of candidate genes whose cardiac expression is modulated by High Sugar and High Fat regimes. Genes encoding core components of key homeostatic pathways - such as 1C-metabolism homeostasis, the Galactose metabolism pathway and metabolites transporters - were identified and characterized as adaptative factors of cardiac function under nutritional stresses. In addition, putative secreted proteins were found dysregulated, highlighting the heart as a secretory organ in hyperglycemia and hyperlipidemia. In particular, we characterized the Fit satiety hormone as a new fly cardiokine, which autonomously modulates the cardiac function and remotely affects feeding behavior. Overall, our study uncovered autonomous and systemic adjustable responses of the heart to nutritional stresses.

## INTRODUCTION

Diabetes mellitus is a chronic metabolic disease characterized by hyperglycemia, affecting 9% of the world’s adult population and responsible for 1.5 million deaths in 2019^1^. The pathophysiological causes and consequences of diabetes are numerous, which makes it a complex disease and a challenge for medicine. Heart diseases, one of the leading causes of death of type 2 diabetics, concern 35% of patients. Diabetic cardiomyopathies are characterized by contractile dysfunctions and structural changes without vascular disorders^2^. The etiology of these specific cardiac dysfunctions remains poorly understood due to the multiple causes of the disease onset. Excessive circulating carbohydrates and lipids lead to insulin resistance in the cardiomyocytes, the trademark of type 2 diabetes (T2D), which concerns 90% of diabetic patients^1^. This induces a lack of glucose import and an abnormal fat accumulation in the cells^2^. Physiological consequences include mitochondria dysfunctions, lipotoxicity, oxidative stress and inflammatory pathways activations, as well as change in stiffness of the cardiac tissue by remodeling of the extracellular matrix (ECM), due to excessive glycation. Cellular dysfunctions can, in extreme cases, lead to cardiomyocyte apoptosis^2^. Resulting cardiac phenotypes are mostly diastolic dysfunctions and increased arrhythmias^2,3^.

*Drosophila*, being the simplest model organism with a beating heart, is a valuable organism for the study of cardiogenesis, cardiac physiology and aging^4–7^. In recent years, it has become a model of choice for the study of metabolic disorders and associated cardiomyopathies^8,9^. The adult cardiac tube is composed of eighty four cardiomyocytes which carry molecular, physiological, and mechanical properties similar to mammalian cardiomyocytes^10^. The contractile cells are in close proximity with a layer of forty pericardial cells, functionally and structurally similar to nephrocytes^11^, with which they closely communicate to sustain their function^10,12^. The physiological equivalence between flies and mammals extends to metabolic perturbations’ incidence on cardiac function. Indeed, pioneering studies have shown that flies fed High Fat (HFD) or High Sugar Diets (HSD) exhibit metabolic disorders and cardiac dysfunctions that are phenotypically reminiscent of mammalian diabetic-induced obesity and diabetic cardiomyopathies, such as abnormal lipidemia, cardiac dysrhythmias and tissue remodeling^9^. Highly conserved metabolic pathways were associated to cardiac perturbations resulting from excess of sugar^13^ or fat^5,14,15^ in the food. It was shown that three weeks HSD feeding led to insulin resistance, body fat accumulation and cardiac fibrosis due to excessive collagen deposits^13^. Moreover, high circulating glucose levels induced activation of the hexosamine pathway in cardiac tissue which negatively impacted its contractile efficacy^13^. Like high sugar, high fat exposure for five days is sufficient to induce obesity and associated cardiac dysfunctions^14^. Indeed, in the fat body and in the heart, high lipids activate the TOR kinase pathway and its lipogenic targets (SREBP and FAS)^14^. More importantly, activated-TOR inhibit the expression of the ATGL/*brummer* (*bmm*) lipase as well as that of its target, the mitochondrial transcriptional co-activator PGC-1/*spargel* (*srl*)^15^. As a consequence, in addition to obesity, this cascade of molecular deregulations induced arrhythmias and excess fat content in the heart, leading to cardiac lipotoxicity, one of the most detrimental consequence of diabetic cardiomyopathies^9,15^.

In this study, we aimed at deciphering the genetic features underlying the cardiac dysfunctions upon nutritional stresses in order to better characterize the onset of diabetic cardiomyopathies. We hypothesized that transcriptomic dysregulations would reflect heart adaptations to this physiological stress and analyzed the cardiac transcriptome of females fed HSD or HFD. Our analysis revealed dysregulation of core homeostatic components such as the one-carbon pathway, the galactose metabolism and metabolites transporters, that we further characterized as key adaptative factors of the heart in response to nutritional stresses. We also observed that both HSD and HFD were accompanied by the dysregulation of a large number of genes encoding secreted proteins, which pointed out the involvement of cardiac myokines (cardiokines) in the adaptation to nutritional stresses. Notably, we identified the *fit* gene, encoding a satiety hormone^16^, as a new fly cardiokine, expressed by cardiomyocytes and pericardial cells, that regulated cardiac function. Importantly, we also demonstrated that cardiac *fit* expression was able to remotely control food intake, further illustrating the adaptability of the heart to nutritional challenge, and its role as a systemic regulator of organismal metabolism and behavior.

## RESULTS

### High-Sugar and High-Fat Diets modified cardiac performance

In order to reveal the early molecular dysregulations induced by high nutrients supply on cardiac function, we investigated how exposure to rich diets affected the cardiac performance. Control females of the strains used thereafter (Figure 1, S1), expressing the cardiac Hand-Gal4 driver (hereinafter referred to as Hand>), were challenged by HSD (10 days) or HFD (7 days in ND + 3 days in HFD) compared to ND (10 days, see Materials & Methods). We evaluated cardiac performance using a noninvasive high throughput assay, based on a heart specific fluorescent transgene, R94C02::tdTomato (attP40) (tdTk), that allows monitoring cardiac activity on intact, anaesthetized flies^17^. Movies of 5s at 330fps were analyzed^18^ to get precise quantification of heart structure and rhythm phenotypes (Figure 1A). Diameters of the cardiac tube at maximum contraction in systole (ESD, End Systolic Diameter) and when fully relaxed in diastole (EDD, End Diastolic Diameter), contractile efficiency (Fractional Shortening, FS; equivalent to the ejection fraction), as well as duration of each heart beat (Heart Period, HP) comprising both contraction (Systolic Interval, SI) and relaxation steps (Diastolic Interval, DI), were evaluated as previously described^7^. Correlated phenotypes, such as arrhythmic events (Arrhythmia Index, AI), the volume of circulating hemolymph in the heart per beat (Stroke Volume, SV) and the hemolymph flow rate (Cardiac Output, CO) were also measured.

**Figure 1.**
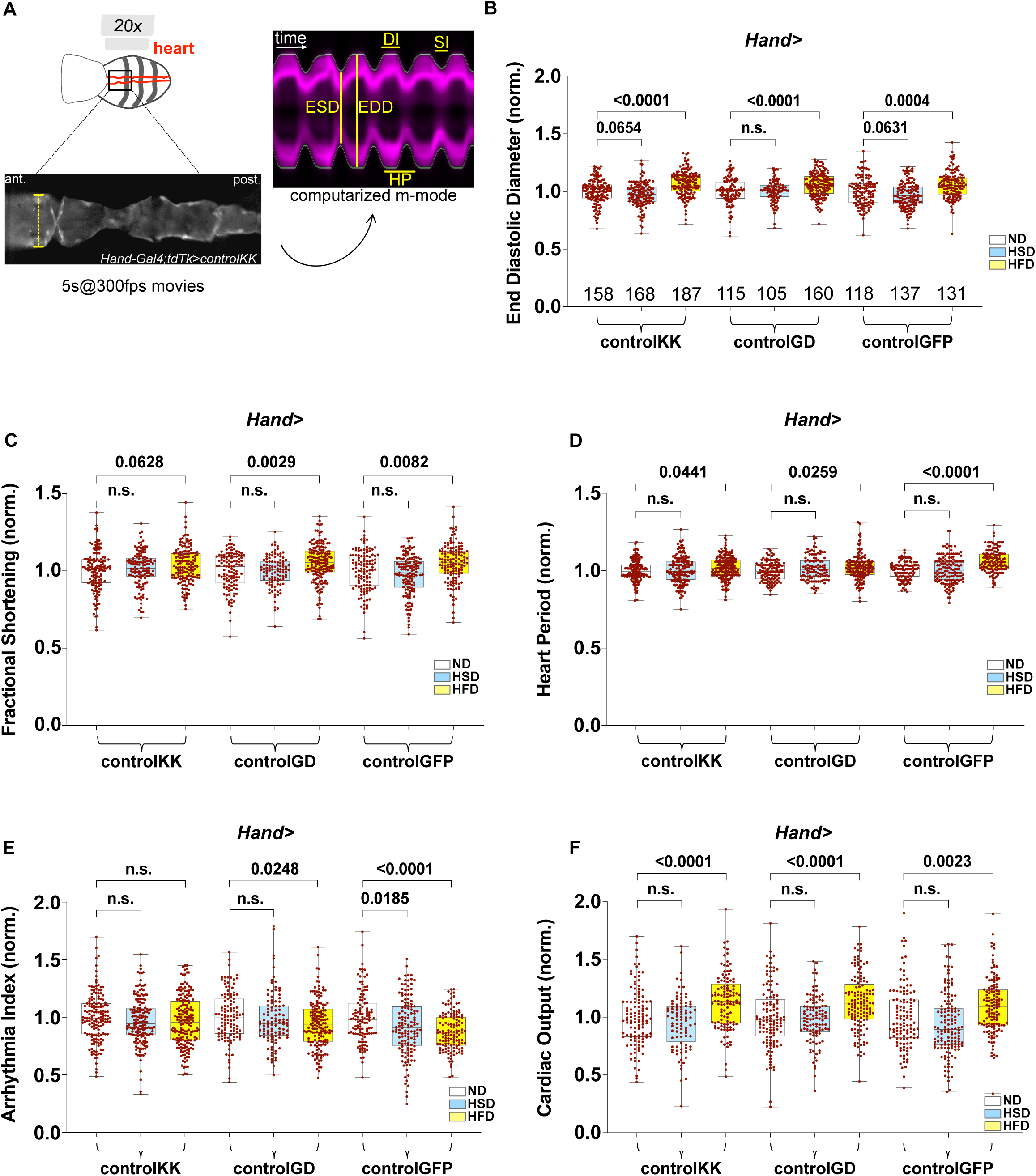
Cardiac performances measured in control flies in response to diets. (A) Live imaging was performed on anesthetized Females (see Methods) expressing the tdTomato (tdTk) reporter in cardiomyocytes. Analysis generated M-Modes (DI=Diastolic Interval; SI=Systolic Interval; HP=Heart Period; EDD=End Diastolic Diameter; ESD=End Systolic Diameter). (B-J) Measurements of cardiac parameters in the 3 types of Hand*>control* flies used in this study following 10days in ND and HSD or 3 days in HFD. Box plots showing normalized values for EDD (B), FS (C), HP (D) and AI (E). Box plots showing normalized values for flux hemolymph estimation in the heart, CO (F). Crosses were performed with Hand>. Phenotypic values of HSD and HFD conditions where normalized to corresponding ND. ND, white plots; HSD, blue plots; HFD, yellow plots. Numbers of individual flies co-evaluated in each diet are indicated. Statistical significance was tested using Kruskal-Wallis with Dunn’s multiple comparisons test. p-values are indicated.

HFD and HSD severely impaired cardiac function in the 3 control genotypes analyzed: heart structure, contractility, rhythmic properties, as well as hemolymph flow were affected (Figure 1, S1). High Fat induced cardiac hypertrophy, as suggested by enlarged diastolic and systolic diameters (EDD, ESD, Figure 1B, S1A) and this was associated to increased Fractional Shortening (FS, Figure 1C). HFD also impacted rhythmicity with increased HP (Figure 1D). Consequently, Stroke Volume (SV, Figure S1D) and Cardiac Output (CO, Figure 1F) were increased. Depending on control strains, reduced AI was observed (Figure 1E). These results indicated that 3 days of HFD is sufficient to trigger deterioration of heart morphology and rate. On the other hand, HSD was associated to milder effects on cardiac physiology. Indeed, it tended to affect several cardiac phenotypes, including EDD and ESD (Figure 1B; S1A), FS (Figure 1C), AI (Figure 1E) as well as hemolymph flux measurements (Figure 1F; S1D), but most of them were not significant and some not consistent between lines. This suggested that cardiac function is less sensitive to glucose-induced stress and may be influenced by genetic background. The discrepancy in observed phenotypes - on both diets - with previous studies^13–15^ could be due to the duration of the nutritional challenge but also to the non-invasive method used here to assess the phenotypes, which does not involve disconnecting the cardiac tube from the central nervous system. Given our objective of studying the initial molecular changes induced, however, we hypothesized that these shorter nutritional stresses would be sufficient and adapted to identify molecular dysregulations.

### Analysis of the fly cardiac transcriptome upon nutritional stress

The same nutritional challenges were used to decipher the molecular perturbations associated to the degradation of the adult cardiac performances described above. Cardiac transcriptome of 10 days female flies was analyzed. Three replicates of 30 hearts were dissected for each diet and sequenced on Illumina NextSeq 500 (Figure 2A). Reads were aligned on dm3 genome annotation and reannotated with dm6 release. After trimming and normalization, 9981 genes were found expressed in cardiac tissue and were subjected to differential expression analysis using both DEseq2_RLE^19^ and edgeR_TMM^20^ (HSD: Figure S2A-C; HFD: Figure S2D-F). Genes found differentially expressed (fold change > 1,5, adjusted p-val < 0,05) with both methods were retained. Compared to ND, respectively 452 (354 down-regulated/80 up-regulated) and 254 (174 down-regulated/80 up-regulated) differentially expressed genes (DEG) were identified in HSD and HFD (Figure 2B-D and Table S1). Detailed analysis of the 110 common DEG between HSD/HFD *versus* ND, showed that 75 had the same pattern (Figure 2E) and 35 the opposite when compared to ND (Figure 2F). Enrichment analyses using PANGEA^21,22^, focused on KEGG annotations, were performed separately on DEG identified in HSD and in HFD *versus* ND. This pointed to carbohydrates and amino-acid metabolism pathways as over-represented among HSD-DEG (Figure 2G-H), whereas lipid, amino-acid and hormone synthesis pathways were identified as over-represented among HFD-DEG (Figure 2I-J). 442 genes were differentially expressed in HSD compared to HFD and exhibited the same patterns of functional enrichment than HSD/HFD *versus* ND (not shown), but they were not analyzed in depth in this context (HSD *versus* HFD, Figure 2D; sugar_*vs_*fat, Table S1). Overall, our data highlighted the modulation of fundamental metabolic processes in the heart upon HSD and HFD (Figure H,J), suggesting a fine tuning of cardiac metabolism in nutritional stresses. We therefore tested whether identified deregulated genes allow adaptative responses of the cardiac function upon these nutritional stresses.

**Figure 2.**
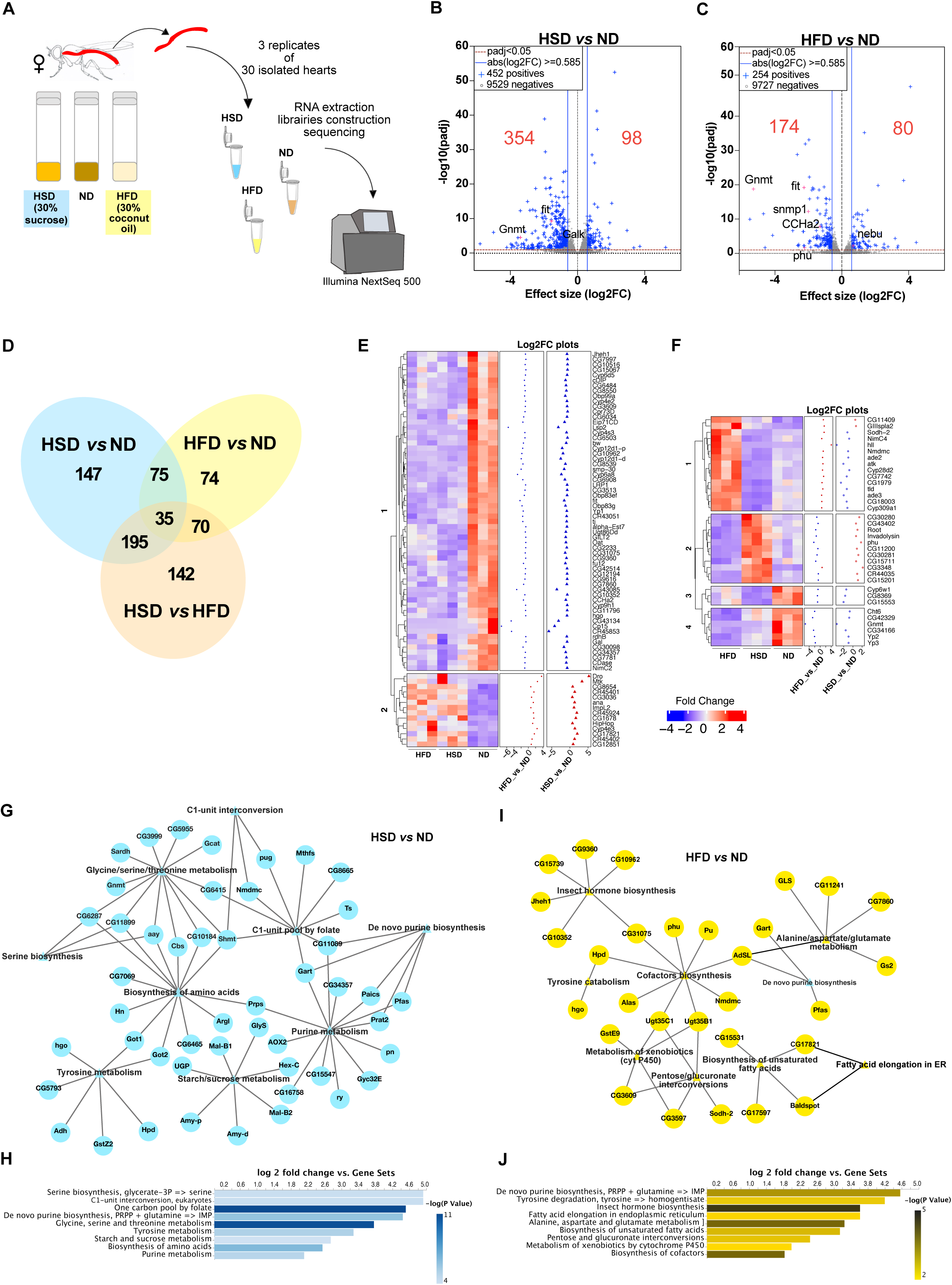
Feeding High fat or high sugar impacts cardiac metabolic regulations. (A) Scheme of the protocol used for batch RNAseq analysis of HFD/HSD-induced cardiac transcriptome modifications. (B,C) Volcano plots showing deregulated genes in HSD and HFD (red numbers). Blue dots indicate genes with an absolute log2FC ≥0.585 and adjusted p-value<0.05. Selected hits are indicated. (D) Venn diagram of DEG identified in the three conditions (padj<0.05). (E,F) Heat maps showing unsupervised clustering analysis and resulting log2FC expression plots for the 75 (E) and the 35 (F) common DEG identified in HSD and HFD stresses (padj<0.05). (G,I) Network graphs obtained with PANGEA enrichment tool for of HSD (G) or HFD (I) DEG. Circle nodes correspond to genes, triangle nodes represent KEGG pathways while edges represent gene to gene association. (H,J) Bar graphs showing log2FC enrichment values for KEGG pathways annotations in PANGEA (padj<0.05).

### High Sugar Diet downregulated 1C-metabolism and Leloir galactose pathways

As outlined above, several genes encoding enzymes associated to 1C-metabolism pathway were found overrepresented in HSD *versus* ND (Figure 2G-H; Figure S2G). 1C-metabolism pathway is a fundamental homeostasis network, which includes several connected pathways involved in methylation (methionine cycle), trans-sulfuration and nucleotide synthesis (folate cycle)^23^. Methionine cycle results in the production of S-adenosylmethionine (SAM), the universal donor of methyl-group in the cell^24^. Among the methyl-transferases that regulates the pool of cellular SAM, GNMT (Glycine-N-methyl transferase) has been involved in the catabolic process of SAM to SAH (S-adenosylhomocysteine), by conversion of Glycine into Sarcosine (N-methylated derivate of Glycine) in the liver in humans^24^ (Figure 3A). SARDH (Sarcosine dehydrogenase) proceeds to the reversal conversion (Figure 3A). Expressed in *Drosophila* cardiac cells (Fly Cell Atlas^25^), *Gnmt* was downregulated in both diets (log2FC: -3.4 in HSD and -5.29 in HFD) while *Sardh* in HSD only (log2FC: -2.05). By RT-qPCR on female hearts, we validate the inhibitory effect of HSD on *Gnmt* and *Sardh* genes expression (Figure S3A). To evaluate the role of 1C metabolism in the heart, the cardiac-specific promoter Hand> was used to perform *Gnmt* and *Sardh* knockdowns in ND, and overexpression of *Gnmt* in ND and HSD. Cardiac performance phenotypes were analyzed using the live-imaging set-up described above (Figure 3B-E). *Gnmt* knockdown (*GnmtKK*) performed in ND fed flies was associated with an increase of HP and with cardiac dilation, with enlarged ESD and EDD (Figure 3C). As a consequence, CO was strongly increased, illustrating an improved hemolymph flux in the heart. Remarkably, while HSD induced an increase in HP in control flies, this was rescued by *Gnmt* overexpression (*Gnmt^WT^*). In addition, while *Gnmt* overexpression did not affected cardiac diameters in HSD, it led to increased contractile efficacy compared to HSD controls (FS, Figure S3F). (Figure 3C). When performed in ND context, *Gnmt* overexpression had no effect on cardiac performance and structure. Overall, our results showed that *Gnmt* is a positive regulator of both cardiac rhythm and contractility, and prevented the HSD-induced decrease in heart rate and improving contractility under this diet. Regarding *Sardh* knockdown, similarly to *Gnmt*, it was associated to increased ESD, EDD and CO in ND (Figure S3C-E). It has already been shown that these two genes similarly influenced the integrity and function of indirect flight muscles (IFMs) in adult flies^26^, indicating that, despite their opposing roles in glycine conversion, they may have the same effect on organ function.

**Figure 3.**
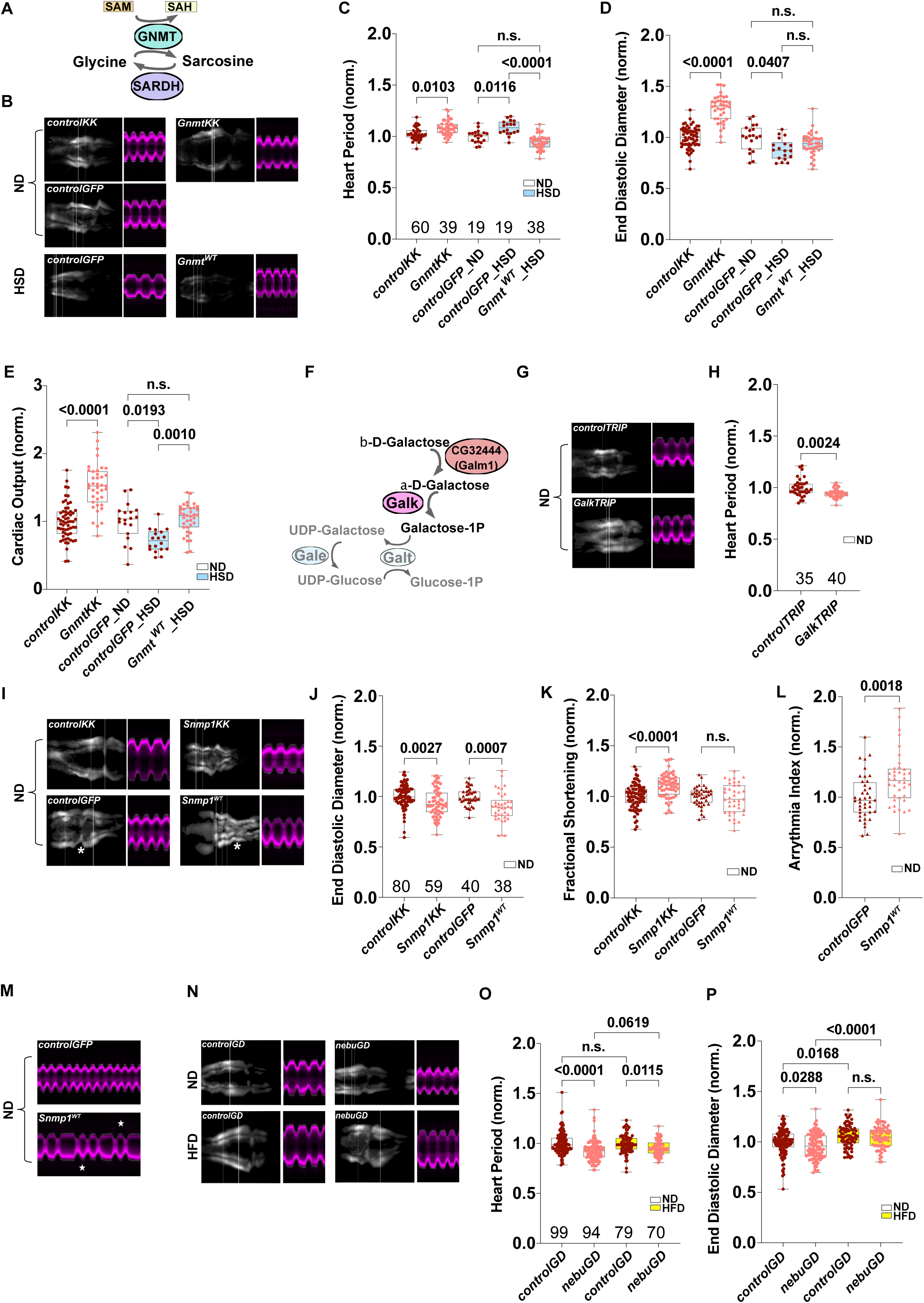
Cardiac function of Glycine metabolism and metabolite transporters. (A) Scheme representing the Glycine cycle, part of the 1C-metabolism pathway. Conversion of Glycine to Sarcosine by GNMT (reversal conversion by SARDH) allows transfer of methyl-group from the universal donor SAM (S-adenosylmethionine) to several cellular components, leading to the formation of S-adenosylhomocysteine (SAH). (B) Z-projection images of heart imaging and generated M-modes from cardiac (Hand>) *Gnmt* knockdown in ND and overexpression in HSD compared to corresponding controls (>*GnmtKK* vs >*controlKK*; >*Gnmt ^WT^* vs >*controlGFP)*. Effect on HP (C), EDD (D) and CO (E) of *Gnmt* knockdown in ND (white box plots) and of *Gnmt* overexpression in HSD (blue box plots) compared to respective controls. (F) Schematic representation of Leloir pathway, comprising enzymes involved in Galactose catabolism. b-Galactose is converted in a-Galactose by CG32444/Galm1, then in Galactose-1P by *Galk*. Following steps lead to Glucose-1P production. Enzymes in grey (Galt and Gale) were not identified in our transcriptomic analysis. (G) Z-projection images of heart imaging and generated M-modes in ND from *Galk* knockdown (*GalkTRIP*) with Hand> and its respective control. (H) HP defects following *GalkTRIP* in ND. (I) Z-projection images of heart imaging and generated M-modes in ND from Hand> driven *Snmp1* knockdown and overexpression in ND compared to respective controls (*>Snmp1KK* vs >*controlKK*; >*Snmp1 ^WT^* vs >*controlGFP)*. Dysfunctional ostia are observed in *Snmp1^WT^* (white star). (J) Defects in EDD and (K) FS upon *Snmp1* knockdown in ND compared to controls. (L) Increased AI induced by *Snmp1* overexpression in ND visualized in corresponding M-modes (M, white stars). (N) Z-projection images of cardiac Hand> driven *nebu* knockdown in ND and HFD compared to corresponding controls (*>nebuGD* vs *controlGD)*. (O) HP and (P) EDD defects following *nebu* KD in ND (white) and HFD (yellow) compared to controls. Crosses were performed with Hand>. Phenotypic values of tested conditions where normalized to corresponding controls. Numbers of individual flies co-evaluated in each diet are presented. Statistical significance was tested using Kruskal-Wallis with Dunn’s multiple comparisons test. p-values are indicated.

HSD also dysregulated enzymes from the Leloir pathway, which are converting α-D-Galactose into Glucose-1P (Figure 3F). *CG32444/Galm1* (*GalM* orthologue) and *Galk* are strongly expressed in cardiac cells^25^ and downregulated in HSD (log2FC: -1.3 and -1.01 respectively) . Fly orthologues of *GalT* and *GalE* were also downregulated, but however did not reached significance (Table S2). By RT-qPCR on female hearts, we validated the inhibitory effect of HSD feeding on *GalK* gene expression (Figure S3A). In ND, *Galk* knockdown was associated to decreased HP and DI (Figure 3G-H, S3I), showing that the effect of *Galk* was to accelerate heart rate by reducing relaxation time. Previous reports have identified *Galk* gene as a suppressor of Calcineurin-induced cardiac dilatation in flies, with no cardiac phenotype by itself when mutated^27^. Our analysis however showed that in normal nutritional context, decreasing *Galk* expression in cardiac cells is sufficient to significantly modulate cardiac rhythm.

### High Fat Diet modulated CD36-scavenger receptor and GLUT8 orthologues

*Snmp1* is strongly downregulated in HFD (log2FC: -2), which we confirmed by RT-qPCR (Figure S3B). *Snmp1* encodes the orthologue of the CD36-scavenger receptor and is strongly expressed in cardiac cells (FlyCellAtlas^25^). *Snmp1* was previously shown to promote Fatty Acids (FA) transport in cells of the larval prothoracic gland and in ovaries, downstream of SUMOylation processes^28^, demonstrating its role in FA metabolism. In human cardiomyocytes, FA and glucose imports both contribute to reach the ATP needs of cardiac contractile cells. FA are the main energetic source of cardiomyocytes, being metabolized by mitochondrial oxidation. Insulin induces translocation to the cardiomyocytes membrane of both FA transporter CD36-scavenger receptor (CD36-SR-B3) and GLUT4 hexose transporter^2^. In insulin-resistance context, GLUT4 mis-translocation and CD36 stabilization result in glucose deficit and FA accumulation in the cells^2^. We observed that, in heart of females fed ND, both *Snmp1* knockdown (*Snmp1KK*) and overexpression (*Snmp1^WT^*) displayed marked structural phenotypes (Figure 3I). EDD and ESD were reduced in both genotypes (Figure 3J; S3J). Modified heart diameters additionally increased FS in *Snmp1KK* (Figure 3K). At the morphological level, abnormal ostia were observed in 100% of *Snmp1^WT^*hearts (Figure 3I, white stars) and were associated to arrhythmias amplification (Figure 3L-M, white stars). We also noticed that excess fat tissue in the abdomen and surrounding the heart was observed in 66% of the imaged *Snmp1KK* females, while it was observed in only 25% of the control flies (Figure S3K). HFD exposure for 3 days induced strong lethality in both *Snmp1KK* and *Snmp1^WT^*, making it difficult to assess the phenotypes with a good statistical power. Nevertheless, our results highlighted a new role of *Snmp1* in autonomous cardiac function with a systemic side-effect on body lipid homeostasis.

HFD also slightly induced the expression of *nebulosa (nebu)* gene (log2FC: 0.6), which trend was confirmed by RT-qPCR (Figure S3B). *nebu* encodes the orthologue of the human SLC2A8/GLUT8 hexose transporter. GLUT8 was recently identified as an insulin-dependent cardiac hexose transporter in mice, whose expression is strongly reduced in diabetes^29^, but its function is not well characterized. Fly gene *nebu* is highly expressed in cardiac cells (FlyCellAtlas^25^). Inhibition of cardiac *nebu* expression, by knockdown (*nebuGD*) in the heart, was performed in ND and in HFD in the same experimental series (Figure 3N-P). In ND, *nebuGD* displayed accelerated heart beats compared to control (revealed by a reduction of HP and DI; Figure 3O; S3L). The same is observed in HFD but HP reduction is weaker (Figure 3O; S3L). in ND, *nebuGD* triggered heart constriction, resulting from a significant reduction of EDD (Figure 3P; S3M), without affecting FS (Figure S3N). However, the hypertrophy induced by HFD was not modified in *nebuGD* condition (Figure 3P; S3M), suggesting that HFD abolished the constrictive effect of *nebuGD*. All these results identified *nebu* as a regulator of heart rate whose constrictive effect in diastole is abrogated by HFD preventing any rescue of the induced-hypertrophy. This could be due to a possible inhibitory regulation of HFD on hexose transporters, as already observed in human cardiomyocytes^2^.

### HSD and HFD revealed the secretory function of the heart

gProfiler^30^ analysis pointed to an enrichment of genes annotated as localized in the extracellular space (GO terms: extracellular region/space) among DEG in both HSD and HFD compared to ND (Figure S4A). Using the BioMart^31^ and SignalP^32^ prediction tools, we identified 123 (in HSD) and 114 (in HFD) cardiac genes encoding putative secreted proteins that are mis-expressed (Figure S4B, Table S3). 51 genes are dysregulated in both diets, including 26 with a characterized human orthologue (Table S4). 22 of these orthologues have been identified in Genome Wide Association Studies (GWAS), exhibiting common variants associated with metabolic or cardiac phenotypes (source “human genetics common metabolic disease knowledge portal”, CMD-KP) (Table S4). Based on homology score (DIOPT^22^) and on the phenotypes associated to their human orthologues, we selected 12 fly genes for specific knockdown in the heart (Figure 4A). *ImpL2*, *Yp1, Yp2, Yp3*, *CG3513*, *Nepl10*, *CG7997*, *phu*, *GILT2*, *CG34357* knockdowns revealed phenotypes reminiscent of the cardiac function related to their orthologues (Figure 4A). Except *phu* and *Invadolysin* (which are inversely dysregulated in HSD and HFD) (Table S1) and *ImpL2* (which is upregulated in both diets), *Yp1-3, CG3513*, *Nepl10*, *CG7997*, *GILT2* and *CG34357* are all downregulated following nutritional stresses. Cardiac knockdown of *Yp2*, *CG3513*, *Nepl10*, *CG7997*, *phu*, *GILT2* and *CG34357*, in ND, were associated to modifications of heart diameters and rate (Figure 4A, seventh column). We observed decreased HP when knocking down *CG34357*, *GILT2*, *CG3513*, *Yp2*, *CG7997* and *Nepl10* (Figure 4B), which resulted from reduction of SI (Figure 4D) and/or DI (Figure S4C). In contrast, both *phu* and *ImpL2* cardiac inhibitions exhibited increased HP (Figure 4B) resulting from extension of the contraction period (SI; Figure 4D). Contractile dysfunction, characterized by a reduced FS, was associated to *Yp2* and *CG7997* knockdown (Figure 4C), consecutive to enlarged ESD (Figure S4D). Reducing *phu* expression also resulted in cardiac hypertrophy (Figure 4E). Despite a high expression in the heart (FlyCellAtlas^25^), knocking down *Invadolysin* led to no detectable cardiac dysfunction (Figure S4F-G as example). Overall, our results strongly suggest that the identified DEG encoding secreted proteins we tested would be putative cardiokines and their human orthologues may be involved in metabolic disorders-associated cardiomyopathies.

**Figure 4.**
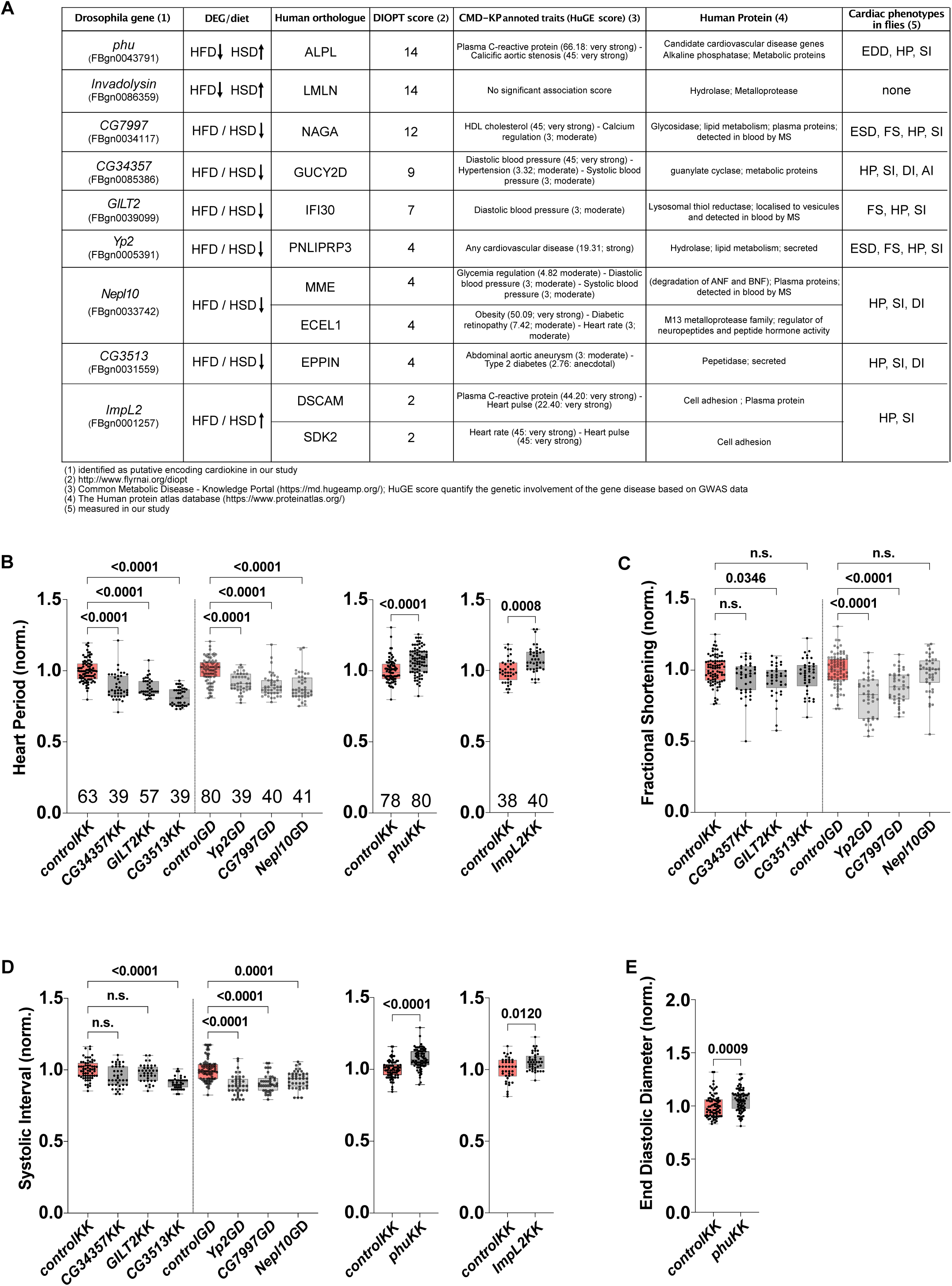
Cardiac function of newly identified cardiokines. (A) Table showing selected putative cardiokine genes identified in HSD and HFD. Fly gene name and Flybase ID (column 1), HSD and HFD dysregulations observed (column 2), best human orthologue (column 3) based on DIOPT score (column 4), HuGE score for diabetes and cardiovascular diseases (quantification of genetic association of genes to diseases based on human GWAS analysis; column 5), protein class and blood detection (Human protein atlas, column 6), identified cardiac phenotypes in fly (column 7). (B) HP, (C) FS and (D) SI modifications observed in Hand> driven KD (grey) compared to respective controls (red). (E) EDD modifications in Hand> driven *phu* KD (grey) compared to control (red). Values where normalized to controls. The number of individuals analyzed in each condition (genotype/diet) is indicated. Statistical significance was tested using Kruskal-Wallis with Dunn’s multiple comparisons test. Significant p-values are indicated. Genotypes: UAS-*CG34357KK*, UAS-*GILT2KK*, UAS-*CG3513KK*, UAS-*Yp2GD*, UAS-*CG7997GD*, UAS- *Nepl10*, UAS-*phuKK*, UAS-*ImpL2KK*.

### Cardiac Fit expression impacted cardiac performance

Among the genes annotated to encode secreted proteins, we selected the gene *fit* for further characterization. Indeed, Fit protein was previously shown to be a satiety hormone^16^, which was an interesting function to investigate as associated to cardiac physiology. In our analysis, the *fit* gene was downregulated both in HSD (log2FC: -1.55) and HFD (log2FC: -2.26) and this was validated by RT-qPCR on dissected hearts (Figure S5A). We then assessed its putative cardiac function by knocking down its expression in ND (using *fitKK* transgene, which reduces its expression by 40%, Figure S5A). In addition to the pan-cardiac driver Hand>, tinCD4-Gal4 (tinCD4>, expressed in cardiomyocytes only^33^) or Dot-Gal4 (Dot>, expressed in pericardial cells only^34^) were used in order to decipher any cell-specificity associated to its function. In all cell types, knocking down *fit* expression was sufficient to reduce HP (Figure 5A-C). Moreover, expressing a modified *fit* isoform, deleted from its signal peptide (*fit^DeltaSP^*), which acts as a dominant negative mutant^16^, resulted in the same HP reduction (Figure 5A). This was mainly due to SI shortening (combined to DI shortening in *fitKK*) (Figure 5D; S5B). Conversely, overexpressing *fit* (*fit^WT^*) was enough to significantly increase HP, in ND and HSD, due to SI lengthening (Figure 5E-F; S5C). However, *fit^WT^* did not affect HP in HFD, which is already increased compared to ND controls (Figure 5E), suggesting a limit to *fit*-induced HP increase. Heart diameters are also dependent on *fit* function. Indeed, inhibiting Fit secretion (*fit^DeltaSP^*) led to heart constriction (reduction of both EDD and ESD; Figure 5G; S5D), despite no significant effect in the *fitKK* knockdown. The reverse phenotypes are induced by *fit* overexpression (Figure 5H, S5E) whatever dietary condition. Indeed, FS was reduced in *fit^DeltaSP^* and augmented in *fit^WT^* (Figure 5I-J), showing that heart diameters and contractility are improved by *fit* in a dosage-dependent manner. AI was decreased with all 3 Gal4 drivers in *fitKK* condition (Figure S5F,H), again suggesting that *fit* is required in all cardiac cells to regulate heart rhythmicity. However, this result was not confirmed in Hand-driven expression of *fit^DeltaSP^* and *fit^WT^* (Figure S5F-G), showing that arrhythmia is also finely regulated by Fit dosage: reduced level (with *fitKK*) promoted arrhythmic events, whereas increased levels did not. Finally, the contrasting phenotypes observed between *fit^DeltaSP^* and *fit^WT^* strongly suggest that cardiac Fit must be secreted in order to exert its functional role in the heart.

**Figure 5.**
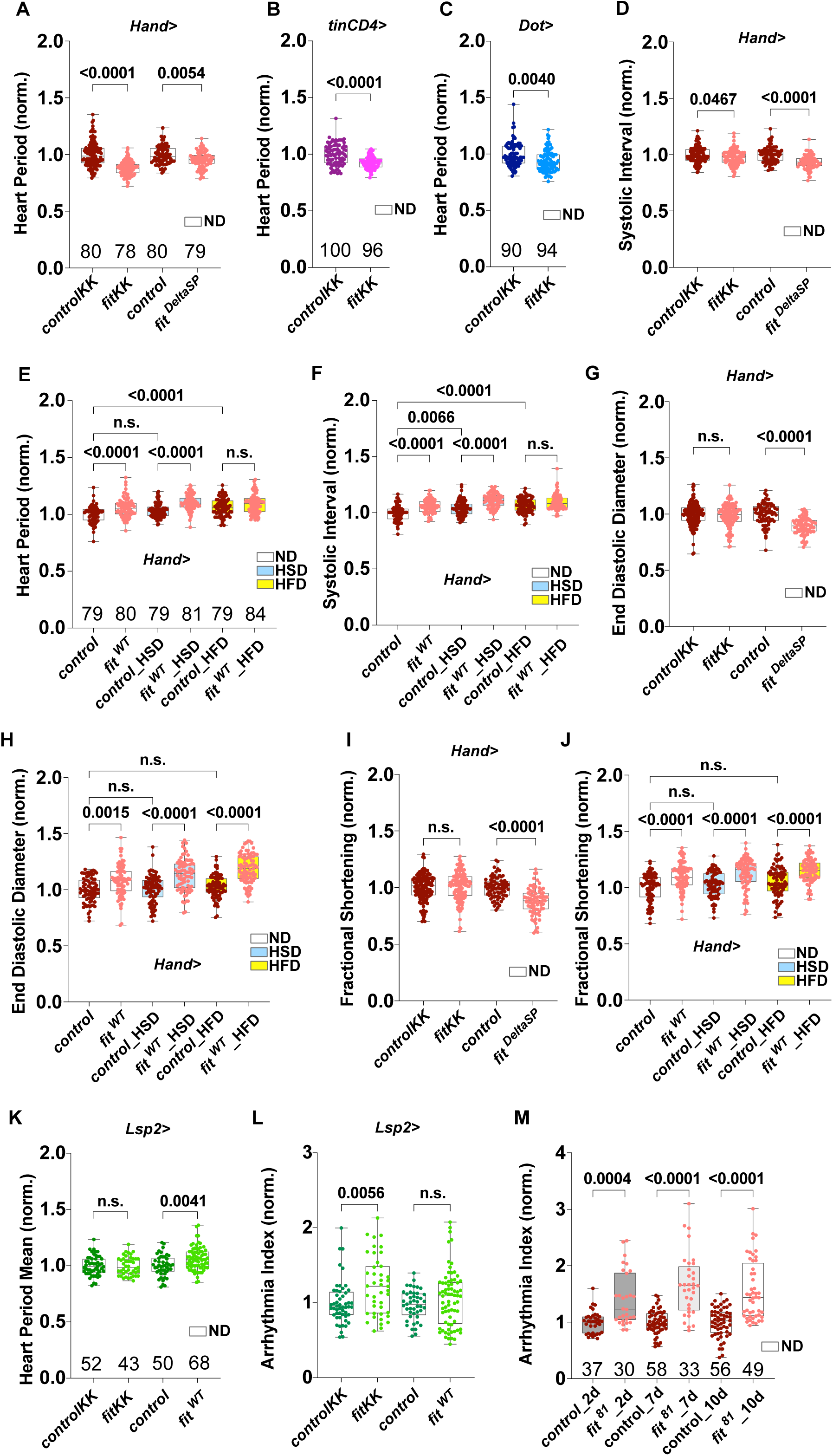
Cardiac Fit is required for cardiac performance. (A) Effect on HP of *fit* knockdown (*fitKK*) and *fit^DeltaSP^* with Hand> in ND. Effect on HP of *fit* knockdown (*fitKK*) in cardiomyocytes (B, *tinCD4>*) and in pericardial cells (C, *Dot>*) compared to respective control flies in ND. (D) Reduced SI observed in *fitKK* and *fit^DeltaSP^* in ND. (E) Effect of *fit* overexpression (*fit^WT^*) on HP and on SI (F) in ND, HSD and HFD compared to corresponding control. (G-H) Effect on EDD of *fitKK* and *fit^DeltaSP^*in ND (G) and of *fit^WT^* in ND, HSD, HFD (H) compared to respective controls. (I-J) Effect on FS of *fitKK* and *fit^DeltaSP^*in ND (I) and of *fit^WT^*in ND, HSD, HFD (J) compared to respective controls. (K) Effect of *fitKK* and *fit^WT^* in ND on HP and AI (L) when expressed in fat body (Lsp2>) compared to respective controls. (M) Effect on AI of mutant *fit*^81^ in females at 2, 7 and 10 days old, compared to controls. Except when mentioned, all crosses were performed with Hand>. Phenotypic values of tested conditions where normalized to corresponding controls. Numbers of individual flies co-evaluated in each diet are presented. Statistical significance was tested using Kruskal-Wallis with Dunn’s multiple comparisons test. p-values are indicated.

We next wondered if Fit secreted from other tissues could have a remote impact on cardiac physiology. We modified *fit* expression in the fat body - a non-cardiac tissue known to express *fit -* with Lsp2-Gal4 (Lsp2>) and analyzed resulting cardiac phenotypic modifications. HP was not modified by adipose *fit* knockdown but was increased by its overexpression (Figure 5K). When focusing on beating phases (Figure S5I-J), increased SI and decreased DI could explain the absence of HP variation in *fitKK,* through a compensatory mechanism. On the reverse, in *fit^WT^*, DI (but not SI) was increased (Figure S5I-J). These results strongly suggested that fat-secreted Fit is also able to remotely control cardiac rhythm. Cardiac dysfunctions, triggered by tissues or organs distinct from the heart itself, were confirmed in a *fit* KO background^16^ *fit*^81^. Indeed, in *fit*^81^, as in Lsp2>*fitKK* (Figure 5L-M), AI is strongly increased. This arrhythmia is observed from the age of 2 days and persisted for more than 10 days (Figure 5M). Even if neither SI, DI nor HP were affected in young *fit*^81^ females, rhythm was already deeply deteriorated in 7-days old ones (Figure S5K-M). At that age, important cardiac morphological defects were concomitantly observed, due to heart wall thickening, which could explain the altered rhythmicity in mutant females (Figure S5N).

Taken together, these results showed that Fit is a regulator of heart rhythm and can also impede on cardiac structure. Cardiac *fit* function is ensured by all cell types and is dependent on its secretion. In addition, *fit* from remote tissues – and in particular the fat body - also impact cardiac functioning.

### Fit, a cardiokine regulating feeding behavior

Fit has been previously identified as a satiety hormone secreted by the fat body^16^. It was shown to regulate food intake by promoting insulin secretion from the IPCs (Insulin Producing Cells), neuroendocrine cells in the *pars intercerebralis* region of the brain^16^. Our analysis identified the *fit* gene as being downregulated under HSD and HFD nutritional stress conditions. We therefore wondered whether, in these diets, cardiac Fit could interfere with insulin levels in the IPCs, fulfilling a new systemic function. First, we tested if cardiac Fit could be secreted into the hemolymph. When expressed in cardiac cells with Hand>, HA-tagged Fit was detected in adult hemolymph extracts (Figure 6A). Then, Hand>*fit^WT^* flies were fed HSD or HFD and Dilp5 immunostaining was performed on brains. Dilp5 is one of the functional orthologues of human insulin in *Drosophila*^35^, produced and secreted by the IPCs. Diet-dependent Dilp5 content in the IPCs reflects its active secretion (low or no staining) or its accumulation (strong staining)^36^. In our control flies (Figure 6B; left panels), Dilp5 could barely be detected in both HSD and HFD compared to its high level detected in ND condition. Of note, due to extreme low signal, some IPCs were impossible to visualize in HFD. In HSD and HFD, Dilp5 amount corresponded to a reduction of more than 50% compared to ND (Figure 6C), and we showed that it was not due to inhibition of *Dilp5* expression by RT-qPCR (Figure S6A). This suggested that both HSD and HFD are sufficient to promote Dilp5 secretion form the IPCs. When *fit* was overexpressed in cardiac cells with Hand>, Dilp5 could not be detected in brains of HSD and HFD fed nor in ND fed flies (Figure 6B, right panels; Figure 6C). In this genotype, whatever the diet, half of the brains had undetectable Dilp5 signal in the IPCs and could not be included in the quantification measurements. RT-qPCR showed that it was not due to the inhibition of *Dilp5* expression in the CNS of these flies (Figure S6A). The reverse experiment, which involved downregulating cardiac *fit* expression, did not show an effect on Dilp5 level in the IPCs or on its expression (Figure S6B-C) in ND. These results corroborate the effect of Fit protein as promoting Insulin secretion from the IPCs, and that Fit can achieve this function from the heart.

**Figure 6.**
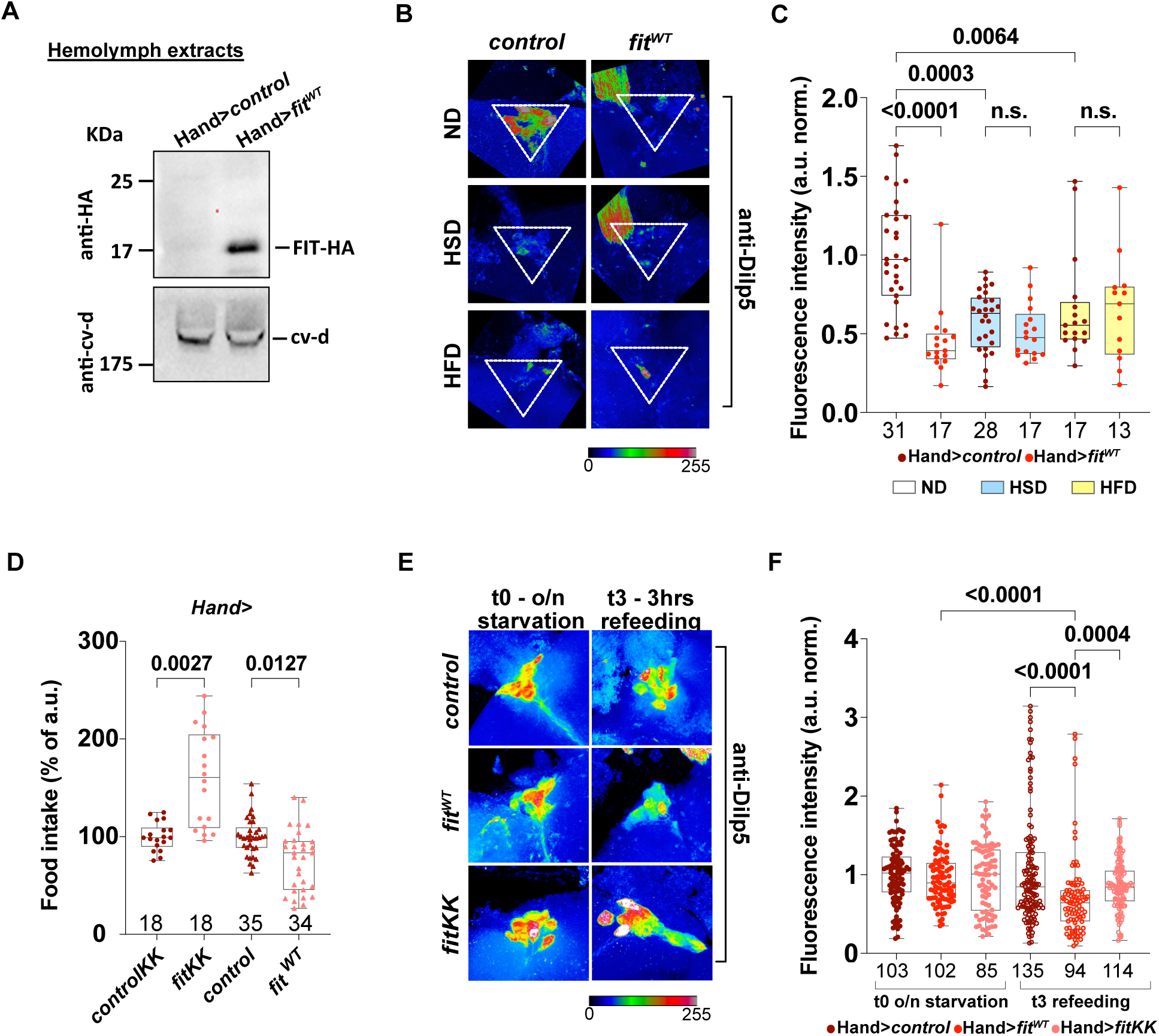
Secreted cardiac Fit promotes Dilp release and regulates food intake. (A) Fit-HA immunodetection in hemolymph extracts from Hand> driven *fit* overexpression (HA-tagged *fit^WT^*) compared to controls. cv-d is a control for hemolymph proteins (n=3 independent experiments). (B) Dilp5 immunostaining in IPCs from dissected adult female brains fed ND, HSD or HFD. Genotypes: Hand>*fit^WT^* (right panels) and controls (left panels). IPCs localization is indicated by white triangles. (C) Fluorescence intensity measured in IPCs from (B) in ND (white boxes), HSD (blue boxes) and HFD (yellow boxes). Values are normalized to ND controls. (D) Quantification of food intake in Hand> driven *fit* knockdown (*fitKK*) and overexpression (*fit^WT^*) females starved overnight and refed 3 hours. Values are normalized to respective controls. Numbers corresponds to averaged values for groups of 10 adults obtained from 3-6 independent experiments. Statistical significance was tested using Kruskal-Wallis with Dunn’s multiple comparisons test. (E) Dilp5 immunostaining in IPCs on dissected adult female brains at t0 (overnight starvation, left panels) and t3 (3hours refeeding ND, right panels). (F) Fluorescence intensity measured in IPCs from to 3 normalized independent experiments related to (E). Genotypes: Hand>*fit^WT^* (light red), *>fitKK* (pink) and controls (dark red). Values are normalized to controls at t0 (t0: points; t3: open circles). In (C, F) Numbers correspond to points recapitulating 3-4 averaged measurements in each IPC analyzed in each condition. Statistical significance was tested using Kruskal-Wallis with Dunn’s multiple comparisons test. Significant p-values are indicated.

To further characterize this remote function, we then evaluated the role of cardiac *fit* in feeding behavior. 1week-old females, fed ND, were starved overnight and then refed ND supplemented with Erioglaucine blue dye for 3hours. Blue food content in gut was measured using lysates from decapitated female bodies (see Methods). As shown in Figure 6D, knocking down *fit* expression with Hand> was sufficient to increase food intake, whereas overexpressing its expression to reduce feeding. We could observe the same kind of trend when *fit* expression was modulated in cardiomyocytes or pericardial cells (Figure S6D), confirming the requirement of *fit* function in all cardiac cell types. These results indicated that Fit secreted from heart cells had a remote effect on a complex behavior such as food intake. In order to correlate these observations with insulin release from the IPCs, we performed Dilp5 immunostaining in adult brains as previously. As already described^36^, a strong Dilp5 accumulation in the IPCs was observed following overnight starvation, in brains from control flies, cardiac *fit* knockdown or overexpression (Figure 6E, left panels). Following 3 hours of refeeding in ND, cardiac *fit* knockdown was insufficient to significantly affect Dilp5 accumulation (Figure 6E-F) but cardiac *fit* overexpressing flies displayed reduced Dilp5 levels in the IPCs compared to refed controls (Figure 6E, right panels, Figure 6F). These observations are not resulting from a change in *Dilp5* (nor *Dilp2*) expression in the adult CNS (Figure S6A; white plots).

These observations globally showed that Fit secreted from the heart was sufficient to remotely modulate Drosophila insulin content in the IPCs, which resulted in the modulation of feeding behavior. All together, these results indicated that the adult heart impeded on systemic metabolic homeostasis and that cardiac *fit* played a key role in this process.

## DISCUSSION

Resulting notably from unhealthy diets, obese diabetics often suffer from cardiomyopathies. In this study, we used *Drosophila* to decipher the genetic and molecular impacts of short High Sugar/High Fat food challenges on cardiac function. Our results revealed key metabolic pathways dysregulated in the heart. 1C-metabolism and Galactose metabolism in HSD, as well as metabolites import transporters in HFD were highlighted. Additionally, we identified numerous putative secreted proteins, suggesting that the heart is able to non-autonomously adapt to diets. Strikingly, we uncovered a cardiac function for the satiety hormone Fit, that impinged locally on cardiac performance and remotely on feeding behavior, highlighting its role as a cardiokine.

Members of purine and folate cycle were downregulated in HSD (Figure S2G), including the two genes *Gnmt* and *Sardh*. Glycine-N-methyl transferase (GNMT) is a key enzyme of the methyl-group donor pool homeostasis. It directly regulates the amount of S-adenosylmethionine (SAM), and its derivative SAH, by conversion of Glycine into Sarcosine. Glycine can be produced back from Sarcosine by SARDH. Knocking down both genes in the heart in ND led to consistent cardiac hypertrophy. *Gnmt* inhibition moreover decreased heart rate and contractile efficacy. Cardiac *Gnmt* over expression in HSD-fed flies was able to prevent the sugar-induced HP increase and improved contractile efficacy, identifying a beneficial role for cardiac *Gnmt* that preserved the heart from the deleterious effects of high sugar diet on rhythmicity and contractile function. These results are in line with known GNMT and SARDH protective effects revealed in time-restricted feeding (TRF) context^35^. TRF was indeed shown to protect heart function from age-dependent degradation^36^ and this was associated, in indirect flight muscles (IFM), to *Gnmt* and *Sardh* upregulation^35^. *Gcat* and *Tdh,* two other genes involved in Glycine regulation in flies, were also slightly down in HSD (Table S1). The cumulative effect of their downregulations with that of cardiac *Gnmt* and *Sardh* upon HSD feeding may lead to a cardiac decrease of Glycine levels and therefore to altered SAM availability. Importantly, in humans, high SAM was associated to obesity^37^, and high circulating SAH levels was shown to be of bad prognosis for CVD^38^. In addition, the circulating level of Glycine was shown to be inversely correlated to CVD risk^39^. This suggests that a tight control of cardiac physiology by 1C- metabolism pathways operate in humans too. Further studies are required to investigate the tight pathophysiological link between circulating sugar, purine cycle and heart dysfunctions.

Sugar stress also unraveled a potential cardiac function of the Leloir pathway. Indeed, several members of this pathway were downregulated under HSD challenge, including (*Galk*) which is involved in the production of Galactose-1P^40^. We showed that *Galk* knock down increased heart rate in ND. The potential contribution of Galactose catabolism to heart physiology in flies is not known, as galactose is not a usual source of nutrient in insects. However, a deficit in Galactose-1P production could compromise subsequent Glucose-1P production which, in turn, could affect cardiac physiology.

We further showed that high fat diet led to a downregulation of *Snmp1* expression, the fly orthologue of the fatty acid (FA) transporter CD36-scavenger, while it slightly upregulated that of the glucose transporter *nebu*, the GLUT8 orthologue. In ND, *Snmp1* knock down in the heart induced heart constriction and impacted contractility. Cardiac inhibition of *Snmp1* was also associated to a marked accumulation of fat in the abdomen and around the heart, suggesting that decreased FA import in the heart is associated to a systemic impact on organismal adiposity. On the other hand, heart specific *Snmp1* overexpression was also associated to cardiac constriction and additionally resulted in strong arrhythmias. These effects are consistent with those observed in humans were CD36 stabilization is deleterious in diabetic cardiomyocytes^2^. In addition, we observed that HFD induced a strong lethality when combined to either inhibition or overexpression of *Snmp1*. Given its known function as a regulator of fat metabolism^28^, our results support *Snmp1* essential function in cardiomyocytes health and a systemic role on lipid homeostasis.

*nebu* knock down lead to cardiac constriction and accelerated heart beats in ND. However, while HFD lead to cardiac hypertrophy in control flies, *nebu* knockdown under hyperlipidemic stress did not modify this cardiac hypertrophy. In humans, T2D insulin resistance is associated with cardiac tissue remodeling, hypertrophy and contractile dysfunction. It was also shown to prevent translocation of the GLUT family glucose transporters at the membrane^2^. In HFD fed flies, a similar mechanism could explain the absence of phenotypes associated with *nebu* inhibition.

Overall, our results highlighted the heart as a metabolic tissue, whose dysfunctions are partly dependent on an autonomous response to nutritional stresses. We also showed that the heart exhibits a non-autonomous response to nutritional stresses. Indeed, 27% of identified DEG encoded putative or known secreted proteins. Remarkably, eight over nine genes encoding such putative secreted proteins were shown to participate in cardiac performance regulation, suggesting they function as paracrine factors to regulate heart performances. Further studies aimed at deciphering their systemic role will be necessary to determine whether they also have a cardiokine function. For instances, *ImpL2* was previously identified as an IGF-binding protein^42,43^ and shown as a cachexia-promoting factor, being secreted from midgut tumoral cells^44,45^. In the cardiac cells under high fat or high sugar stress, secreted ImpL2 could also, as a cardiokine, affect peripheral tissues physiology. Identification of biomarkers for cardiac stress in patients is an important field of research for early diagnosis of diabetic cardiomyopathies. Our study thus provides a promising repertoire of cardiokines in fly that may prompt research on their orthologues in mammals.

In order to improve our understanding of the secretory properties of the heart, we focused on the *fit* gene, whose cardiac expression is downregulated in response to both HSD and HFD. Fit has been previously characterized as a fat-secreted satiety hormone in response to protein feeding^17^, that also regulate sugar intake for sustained egg laying in females^46^. In ND, inhibiting its expression in the heart increased heart rate, the reverse being observed following its overexpression. Additionally, *fit* impacted cardiac structure, leading to hypertrophy when overexpressed in HSD and HFD, consequently affecting heart contractility. We showed that cardiac Fit was secreted in the hemolymph and that Fit deleted of its signal peptide acted as a dominant negative allele that did modify heart performances. This demonstrated that cardiac expressed Fit acts in an autocrine manner to control cardiac function. In addition, when *fit* expression was inhibited or overexpressed in fat tissue, contractile and rhythmic cardiac degradations were observed. Arrhythmic events increase was also observed both in *fit* KO females and following *fit* knockdown in the fat body. Overall, these results established both an autocrine and a paracrine function of Fit as a crucial regulator of cardiac performance. Functional characterization of Fit expressed from the heart also showed that it remotely impacts insulin levels, promoting its secretion from the IPCs when overexpressed in ND. We hypothesized that it would modify food intake accordingly. Indeed, feeding amount was inversely correlated to the level of cardiac *fit* expression. These results showed that secreted cardiac Fit could signal to the adult brain to regulate feeding behavior via controlling insulin release form the IPCs. By acting both autonomously within the heart and remotely on insulin release and food intake, Fit possesses all the attributes of a cardiokine. Studies in mammals identified several cardiokines characterized in response to cardiomyocyte stresses and cardiac remodeling processes^47^, some of which remotely controlling metabolic homeostasis^48,49^. In flies, several studies identified proteins secreted from cardiac cells that act as regulators of lipid homeostasis and organismal obesity^50–52^. Here we identify a cardiac secreted protein which not only impinges on cardiac performance but also affects food intake. Our results therefore highlight the heart as an active metabolic organ and a key physiological regulator able to remotely impact on feeding behavior.

## MATERIALS AND METHODS

### Fly husbandry and stocks

All lines were reared in incubator at 18°C for husbandry on reference food (ND, normal diet) and in incubators (light/dark cycle of 12/12h, humidity of 50%), at 23°C or 25°C for experiments in selected food as indicated. Animals were reared on normal diet (ND) at 25°C (or 23°C) containing per liter: 10g agar, 70g corn flour, 70g inactivated yeast extract and 3,75g Moldex (in Ethanol). High sugar diet (HSD) corresponds to the same food supplemented with 30% sucrose and high fat diet (HFD) with 30% coconut oil. For the purpose of experiments, flies were kept on ND or HSD at 25°C for 7 days then transferred at 23°C on ND, HSD or HFD for 3 more days.

The following lines were obtained from the BDSC: w^1118^ (BDSC#3605; RRID:BDSC_3605); *Dot-GAL4* (BDSC#67608; RRID:BDSC_67608); *tinC-Gal4.Delta4* (BDSC#92965; RRID:BDSC_92965); *UAS-GFP.Val10* (*controlGFP*; BDSC#35786; RRID:BDSC_35786); UAS-*Snmp1^WT^* (BDSC#25044; RRID:BDSC_25044).

The following UAS-RNAi lines were obtained from the VDRC (RRID:SCR_013805): control lines #60100 (VDRC#60100, for KK lines) and #60000 (VDRC#60000, for GD lines); *UAS-GnmtKK* (VDRC#110623); *UAS-SardhKK* (VDRC#108873); *UAS-Snmp1KK* (VDRC#104210); *UAS-CG10960GD* (VDRC#8359); *UAS.CG34357KK* (VDRC#105185); *UAS-GILT2KK* (VDRC#104414); *UAS-CG3513KK* (VDRC#107668); *UAS-ImpL2KK* (VDRC#106543); *UAS-phuKK* (VDRC#100052); *UAS-Yp2GD* (VDRC#50156); *UAS-CG7997GD* (VDRC#16840); *UAS-Nepl10GD* (VDRC#7736); *UAS-InvadolysinGD* (VDRC#16416); *UAS-fitKK* (VDRC#109482); *UAS-fitGD* (VDRC#14434).

The following lines were obtained from FlyORF : UAS-*Gnmt^WT^* (FlyORF#F04086); UAS-*fit*.3XHA (FlyORF#F002469; RRID: FlyBase_FBst0501830) called UAS-*fit^WT^* in this study.

Other lines: *Hand*^4.2^*-Gal4* (gift from Dr Achim Paululat, University of Osnabrück, Germany); *Hand*^4.2^*-GAL4;R94C02::tdTomato* and *tinC-Gal4.Delta4;R94C02::tdTomato* (gift from Dr Karen Ocorr, SBP Medical Discovery Institute, San Diego, USA); *Dot-Gal4;R94C02::tdTomato* (this study); *Lsp2-Gal4* (gift from Dr Pierre Léopold, Institut Curie, Paris, France); *Lsp2-Gal4;R94C02::tdTomato* (this study); UAS-*fit ^DeltaSP^* and *fit*^81^ (from ^17^ were kindly given by Carla Margulies, Biomedical Center Munich, Ludwig Maximilians University, Germany).

### RNA extraction and sequencing

Progeny from w1118 line was obtained from calibrated tubes (20 females and 5 males per tube) at 25°C. Flies were reared on ND and HSD at 25°C for 7 days (20 females and 5 males per tube; males removed after 2 days; tubes are changed every 2 days). Mated females were then transferred at 23°C in order to get one tube of 20 females on ND, HSD and HFD per date. Three independent samples were collected at three different days for each food condition. For each sample, hearts of 30 females of each condition were dissected and immediately put in a collection tube containing 100ul of TRIzol (ThermoFisher Scientific, #15596026). RNA extraction was performed as described in ^53^. RNA quantification was performed on Nanodrop (DS-11FX, DeNoviX Inc.) and Qubit (Invitrogen). The RNA-sequencing was performed on three independent samples (containing each >500ng) for each food condition. cDNA librairies were constructed and paired-end 50/30 Illumina sequencing realized (NextSeq-500; TAGC/TGML core facility, https://tagc.univ-amu.fr/en/resources/transcriptomic-genomic-platform-marseille-luminy).

Quality control of the data was performed with FastQC v0.11.7 (available online http://www.bioinformatics.babraham.ac.uk/projects/fastqc/). The following tools were used to analyse the raw data: trimming was realized with Sickle (v1.33; https://github.com/najoshi/sickle; Joshi, N.A.; Fass, J.N., 2011); mapping on Dm3 genome release with Subread aligner (subread-align with subread v1.6.1^55^); Read counts with featureCounts (with subread v1.6.1^56^). Normalization of reads was implemented in Dseq2 and edgeR Bioconductor packages (RLE for DESeq2 and TMM for edgeR) ^20,21^. Differentially Expressed Genes (DEG) was calculated in three conditions HSD *versus* ND, HFD *versus* ND and HSD *versus* HFD. Genes with a differential expression ≥1.5 in fold change and with a p-val <0.05 in both methods were selected.

Clustering of the datasets was performed using the unsupervised K-means learning algorithm and heatmaps realized with “pheatmap” v1.0.12 (https://rdocumentation.org/packages/pheatmap/versions/1.0.12 ) and “ComplexHeatmap”v 2.12.0 (available from the Bioconductor project: http://www.bioconductor.org/packages/devel/bioc/html/ComplexHeatmap.html Gu et al., 2016) packages in R/Rstudio (RStudio Team (2022). RStudio: Integrated Development Environment for R. RStudio, PBC, Boston, MA; URL: http://www.rstudio.com/ ); R project version 3.5.1 (RRID:SCR_001905). Validation of gene_id was performed with FlyBase ID validator tool (http://flybase.org/convert/id) in Dm6 genome annotation (Flybase: RRID:SCR_006549).

### Functional annotation and sequence analysis

Functional annotation was performed using g:Profiler public resource (https://biit.cs.ut.ee/gprofiler/gost, RRID:SCR_006809) ^30^. The complete list of TRUE significative DEG for each nutritional context was submitted as a query for “only annotated genes”; Benjamini-Hochberg FDR was chosen for the significance threshold filtering of enrichment terms (p=0.05) and search was run “Gene Ontology” databases (GO MP, GO BP, GO CC).

Prediction of signal peptides in the candidate genes sequences was done with BioMart on Ensembl project (https://www.ensembl.org/biomart/martview/, Ensembl RRID:SCR_002344). Query was performed on the BDGP6.2 *Drosophila* gene dataset on complete list of DEG for each condition (using the gene stable ID FBgnxxxx as a filter) and cleavage site (https://services.healthtech.dtu.dk/services/SignalP-5.0/) was evaluated as a result. Enrichment map of KEGG pathways were realized using PANGEA^22^ v1 beta 1 (https://www.flyrnai.org/tools/pangea/web/home/7227) with “KEGG pathways” as a proxy .

### Fly heart live recording

Heart wall videos were performed on anaesthetized females expressing the *R94C02::tdTomato* transgene. 10 days old adult females are anaesthetized with FlyNap (Sordalab, #FLYNAP). Flies are then glued on a coverslip, on the dorsal side, in droplets of UV-curing glue (Norland Optical, #NOA61). After polymerization, the coverslip is fixed on a small petri dish, the dorsal side of the flies facing the 20x dry objective on a Zeiss Axioplan microscope. Cardiac activity is recorded in the A2-A3 abdominal region of the heart tube with a HD/high-speed camera (OrcaFlash4.0 digital CMOS camera, Hamamatsu Photonics). A 5sec movie at 300 frames/sec is recorded using HCI imaging software (Hamamatsu Photonics). Movies are analyzed with a designed R/Rstudio script (https://github.com/gvogler/FlyHearts-tdtK-Rscripts)^19^, allowing the generation of M-modes and the quantification of different cardiac phenotypes. Cardiac parameters were calculated as described previously^18^: Diastolic interval (DI) is the duration (in seconds) for each heart relaxation phase (diastole). Systolic interval (SI) is the duration (in seconds) of the contraction phase of the heart. Heart period (HP) is the time between the two consecutive diastoles (HP=DI+SI, in seconds) and heart rate is calculated as HR=1/HP (in Hertz). Arrhythmia index is the standard deviation of the HP mean for each fly normalized to the median HP (AI=std-dev (HP)/ mean HP). Fractional shortening is the percentage of diameter change at each contraction and is calculated as FS=EDD-ESD/EDD; Stroke Volume (in picoliters) is SV= π( 1/2(EDD)^2^) – π (1/2(ESD)^2^) and Cardiac Output (in picoliters/seconds) corresponds to CO=SV/HP. Values were normalized to controls/date of experiment (https://gitlab.com/krifa_sallouha/TDTK_Data_AnalysisScripts).

### RT-qPCR

Female adult hearts (Figure S3A-B, S5A) or heads (Figure S6A) were dissected in artificial hemolymph solution^58^, then frozen in liquid nitrogen. Total RNA was extracted using Qiagen miRNeasy Mini Kit (#217004, for hearts) or with RNeasy Plus Mini Kit (#74134, for heads) according to the manufacturer protocol. RNA samples were reverse-transcribed using SuperScript™ VILO™ Master Mix (Invitrogen #11755*) and the generated cDNA used for RT-PCR (QuantStudio 6 Flex, Applied Biosystem) using PowerSYBRGreen PCR mastermix (Applied Biosystem #4367659) as previously described ^59^. Three biological replicates (10-15 hearts or 15 heads) were collected for each sample and technical triplicate was conducted for each. (Primers sequences are provided in Supplemental Material and Methods). Analysis of the data were performed using the software provided with the equipment.

### Feeding behavior

10 days old adult females were starved overnight on PBS/Agarose vials. They were then challenged for 3 hours on tubes containing ND supplemented with 0.5% Erioglaucine disodium salt (Sigma Aldrich, #861146). Flies were frozen as group of 10/1.5ml Eppendorf tube until OD measurements at 629nm as described previously ^60^.

### Western-blotting

Hemolymph samples from 20 females/lane, reared on ND, were extracted as described in ^61^, and prepared for Western Blotting. Hand>*control* and Hand>*fit^WT^* were tested. The experiment was performed 3 times independently. cv-d was used as a normalizer for hemolymph proteins ^62^. Proteins were resolved by SDS-PAGE, using 4-12% gradient gels (NuPage Novex Gel, InVitrogen #NP0335). Primary antibodies used were: rat anti-HA (1:1000, Merck/Roche, #ROAHAHA, RRID:AB_2687407) and guinea-pig anti-Cv-d (1:2000 ^62^, gift from Eaton lab). Membranes were washed in PBS tween 0,1% and incubated with secondary antibodies in this buffer for 2 hrs at room temperature. The following HRP conjugate secondary antibodies were used (Thermo Fisher Scientific): Goat anti-rat (1:2500, #A-10549, RRID:AB_10561556) and anti-guinea pig (1:2500, #A-18769, RRID:AB_2535546). Chemiluminescence was observed using the ECL plus detection substrate kit (Thermo Fisher Scientific #32134). Images were generated using Fiji^63^ (ImageJ version 2.0.0-RC-69/1.52n; RRID:SCR_002285).

### Immunostaining

Adult brains were dissected in cold PBS on ice, fixed in 3,7% methanol-free formaldehyde (Polysciences Inc., #18814-10) for 25 min, washed several times in PBS + 0,1% Triton X-100 (PBT) before 2 hrs blocking in PBT containing 10% BSA. Primary antibody, Rabbit anti-Dilp5 (1:800, generated in Leopold’s lab)^64^ was incubated overnight at 4°C. Alexa conjugated secondary antibody (Alexa Fluor 546 goat anti-rabbit: #A-11035, RRID:AB_143051) was incubated 2 hrs at room temperature. After several washes, brains were mounted in Vectashield (Eurobio scientific, #H1000-10). Fluorescence images were acquired using a Zeiss LSM780 (63x immersion objective) and processed with Fiji^63^ (ImageJ version 2.0.0-RC-69/1.52n; RRID:SCR_002285).

### Antibody staining quantification

For quantification, stacks were taken for the whole IPCs cluster in each brain, from top to bottom limit of the staining. Slices of the whole stack were used to evaluate the fluorescence intensity using Fiji^63^ (StagReg, Time Series Analyser V3.0 and Roi Manager plugins). For each cluster, 3-4 individual neurons per cluster were measured as described previously^65^.

### Statistical analysis and Figures design

Statistical analyses and graphical representations were performed with Graphpad Prism V10 (© 2024 GraphPad Software; RRID:SCR_002798). The statistical test used for each experiment and the p-values are indicated in the corresponding Figure and Figure legends. Fiji (ImageJ version 2.0.0-RC-69/1.52n; RRID:SCR_002285) was used for Image treatment from microscopy and Western Blotting. Affinity Designer v.1.10.8 (Serif, Europe; RRID:SCR_016952) was used for Figures elaboration.

## Data availability

RNA-seq raw data have been deposited in the NCBI Gene Expression Omnibus (accession no.GSE235313) (RRID:SCR_005012).

## Acknowledgments

We thank Dr Achim Paululat, Dr Karen Ocorr, Dr Carla Margulies and Dr Pierre Léopold for fly stocks and reagents. We thank Dr Pauline Brochet for her kind help with GEO submission, Frederic Gallardo and Tahagan Titus for their technical assistance in fly food cooking. Undergraduate students Charis Aubert, Alice Corbet and Loïc Crespo participated to the project during their master internship. We thank the Bloomington Stock Center and the Vienna *Drosophila* RNAi Center for fly stocks. We thank the TGML sequencing platform and E. Castellani from the IBDM Imaging Facility and France Bio-Imaging infrastructure - ANR-10 INBS-04-01.

## Funding

Funding is from Amidex (grant n°AMX-21-PEP-020 to N.A.), ANR (grant n°ANR-22-CE17-0051-03 to N.A.), Fondation de France (grant n°00071034 to L.P.), Inserm and Aix-Marseille University (AMU).

## Contributions

Conceptualization: L.P., N.A., Data Curation: L.K.C., L.K., Formal Analysis: L.K.C, L.K., N.A., Methodology: N.A., S.K., L.P., Experimental Investigation: N.A., L.K., M.T., C.A., L.C., A.C., L.R., Project Administration: N.A., L.P., L.R., Supervision; N.A., Validation: N.A., Visualization: N.A., Writing- Original Draft Preparation: N.A., L.P., Writing- Review & Editing: N.A.

---

**Figure S1. Cardiac performances measured in control flies in response to diets (related to Figure 1).** Measurements of cardiac parameters in the 3 types of Hand*>control* flies used in this study following 10days in ND and HSD or 3 days in HFD. Box plots showing normalized values for ESD (A), DI (B), SI (C) and SV (D). ND, white plots; HSD, blue plots; HFD, yellow plots. Numbers corresponds to the individuals co-evaluated in each diet. Statistical significance was tested using Kruskal-Wallis with Dunn’s multiple comparisons test. Significant p-values are indicated.

**Figure S2. Comparative analysis of the transcriptome with DEseq2 and edgeR_TMM (related to Figure 2).** (A-C) Compared analysis for HSD *versus* ND results. (D-F) Compared analysis for HFD *versus* ND results. (A,D) Venn diagrams of selected genes (padj. <0.05, FC>=1.5). After filtering, 9981 genes were analyzed with each method. Sugar or Fat hits in yellow and normal genes in blue. (B,E) Comparison of genes according to log2FC (left panels) or padj. value (right panels) with each method. (C,F) Volcano plots showing the repartition of the total genes in each condition determined in DEseq2 *versus* edgeR_TMM. padj<0.05, abs(log2FC)>=0.585, blue crosses indicate significant hits *versus* grey n.s. (G) Volcano plot highlighting members of the 1C-metabolism pathway downregulated in HSD *versus* ND condition.

**Figure S3 (related to Figure 3).** (A-B) RT-qPCR profiles of *Gnmt, Sardh* and *GalK* expression in HSD vs ND; (B) RT-qPCR profiles of *Snmp1* and *nebu* expression in HFD vs ND. Fold Changes are normalized to *RP49*. Statistical significance was tested unpaired t-test with Welch’s correction. (C) Effect on ESD of *Gnmt* knockdown in ND (white box plots) and of *Gnmt* overexpression in HSD (blue box plots) compared to respective controls. (D-E) Effect of driven *Sardh* knockdown (*SardhKK*) on EDD (E) and ESD (F) compared to *>controlKK* in ND. (F) Effect on FS of *Gnmt* knockdown in ND (white box plots) and of *Gnmt* overexpression in HSD (blue box plots) compared to respective controls. (G-H) Effect of *Sardh* knockdown (*SardhKK*) on FS (E) and CO (F) compared to *>controlKK* in ND. (I) Effect of *GalK* knockdown (*GalkTRIP*) on DI compared to control in ND. (J) Effect of *Snmp1* knockdown (*Snmp1KK*) and overexpression (*Snmp1^WT^*) on ESD compared to respective controls in ND. (K) Image capture from representative movies showing abdominal fat accumulation (arrowhead) in Hand> driven *Snmp1KK* and control. (L-O) Effect of *nebu* knockdown (*nebuGD*) on DI (L), ESD (M), FS (N) and AI (O) compared to controls in ND (white box plots) and in HFD (yellow box plots). Crosses are performed with Hand>. Phenotypic values of tested conditions where normalized to corresponding controls. Numbers of individual flies co-evaluated in each diet are presented. Statistical significance was tested using Kruskal-Wallis with Dunn’s multiple comparisons test. p-values are indicated.

**Figure S4 (related to Figure 4).** (A) gProfiler graphical representation of enrichment analyses for cellular components GO terms on total DEG identified in HFD (yellow) and HSD (blue) *versus* ND. (B) Venn diagram of DEG identified with Biomart/SignalP 5.0 in HSD and HFD *versus* ND (padj<0.05). (C) DI and (D) ESD modifications in Hand> driven KD (grey) compared to respective controls (red). (E) Effect of Hand> driven *Invadolysin* KD on HP and (F) FS compared to control. Values where normalized to controls. The number of individuals analyzed in each condition (genotype/diet) is indicated. Statistical significance was tested using Kruskal-Wallis with Dunn’s multiple comparisons test. Significant p-values are indicated.

Genotypes: UAS-*CG34357KK*, UAS-*GILT2KK*, UAS-*CG3513KK*, UAS-*Yp2GD*, UAS-*CG7997GD*, UAS-*Nepl10*, UAS-*InvadolysinGD*.

**Figure S5 (related to Figure 5).** (A) RT-qPCR profiles of *fit* expression in HSD or HFD vs ND, and in Hand> driven *fitKK* knockdown vs *>controlKK* in ND. Fold Changes are normalized to *RP49*. Statistical significance was tested unpaired t-test with Welch’s correction. (B-C) Effect on DI of *fitKK* and *fit^DeltaSP^*in ND (B) and of *fit^WT^*in ND, HSD, HFD (C) compared to respective controls. (D-E) Effect on ESD of *fitKK* and *fit^DeltaSP^* in ND (D) and of *fit^WT^* in ND, HSD, HFD (E) compared to respective controls. (F-G) Effect on AI of *fitKK* and *fit^DeltaSP^*in ND (F) and of *fit^WT^* in ND, HSD, HFD (G) compared to respective controls. (H) Effect on AI of *fit* knockdown (*fitKK*) in cardiomyocytes (B, *tinCD4>*) and in pericardial cells (C, *Dot>*) compared to respective control flies in ND. (I-J) Effect of *fitKK* and *fit^WT^* in ND on SI (I) and DI (J) when expressed in fat body (Lsp2>) compared to respective controls. (K-M) Effect of mutant *fit*^81^ in females at 2, 7 and 10 days old, on SI (K), DI (L) and HP (M) compared to controls. (N) Image capture from representative movies showing thickening of the heart wall and obstructed heart lumen in *fit*^81^ mutant females compared to control at 7 days-old.

Except when mentioned, all crosses were performed with Hand>. Phenotypic values of tested conditions where normalized to corresponding controls. Numbers of individual flies co-evaluated in each diet are presented. Statistical significance was tested using Kruskal-Wallis with Dunn’s multiple comparisons test. Significant p-values are indicated.

**Figure S6 (related to Figure 6).** RT-qPCR profiles of *Dilp5* and *Dilp2* expression in (A) ND, HSD and HFD in Hand>*fit^WT^* (dashed column bars) compared to controls (plain column bars) and in (B) Hand>*fitKK* (checkered column bars) compared to controls (plain column bars). Fold Changes are normalized to *RP49*. Statistical significance was tested unpaired t-test with Welch’s correction. (C) Fluorescence intensity of Dilp5 immunostaining measured in IPCs from dissected adult female brains fed ND. Genotypes: Hand>*fitKK* and Hand>control. Values are normalized to controls. Statistical significance was tested using Kruskal-Wallis with Dunn’s multiple comparisons test. (D) Quantification of food intake tinCD4> or Dot> driven *fit* knockdown (*fitKK*) and overexpression (*fit^WT^*) females starved overnight and refed 3 hours. Values are normalized to respective controls. Numbers corresponds to averaged values for groups of 10 adults obtained from 3-6 independent experiments. Statistical significance was tested using Kruskal-Wallis with Dunn’s multiple comparisons test.

## Notes

### Competing Interest Statement

The authors have declared no competing interest.

### Summary of Updates

Full revised text and figures supported by new analysis and experiments performed

