## Supplementary figures and images for "Metabolism fine tuning and cardiokines secretion represent adaptative responses of the heart to High Fat and High Sugar Diets in flies"

### Figure S1

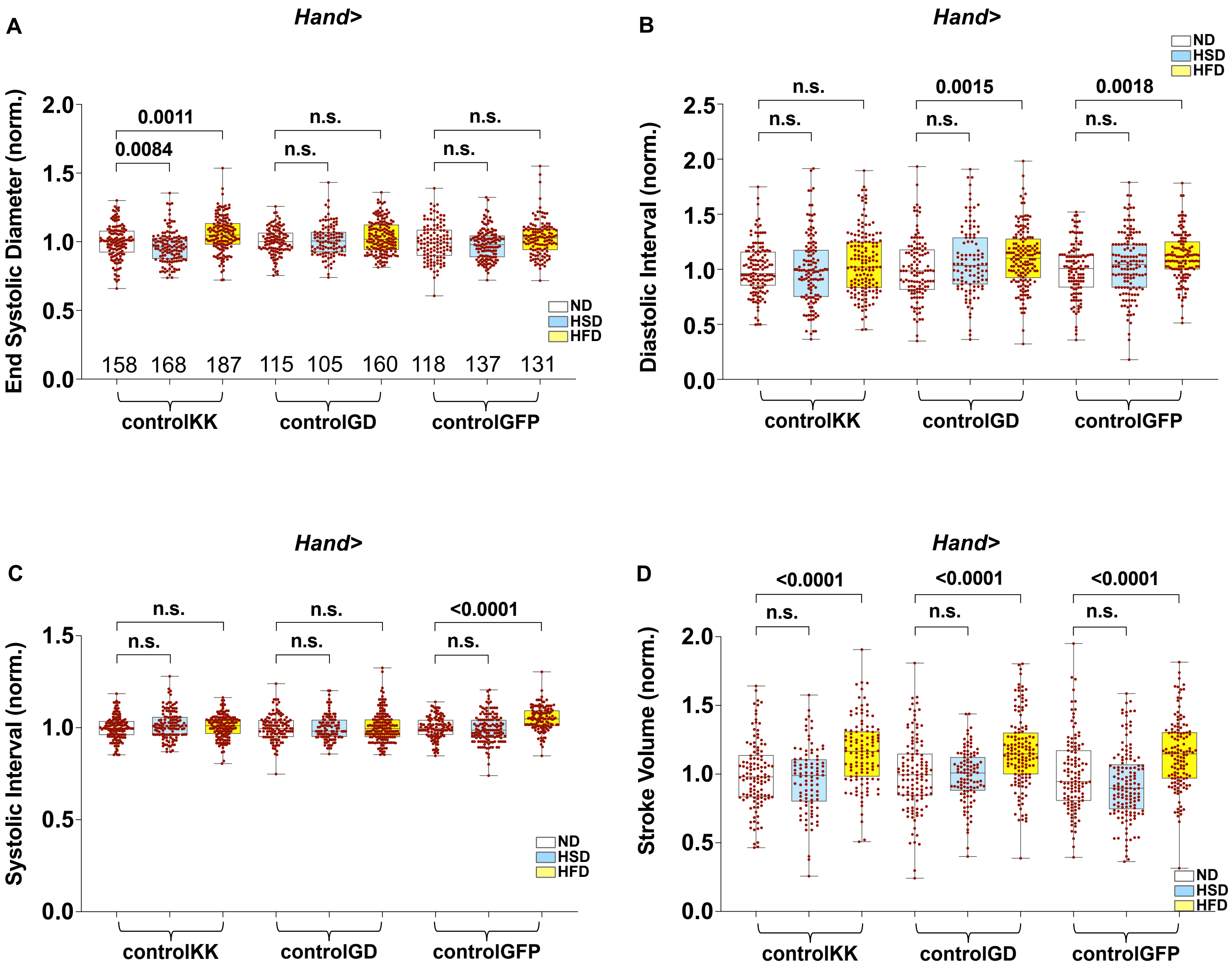

### Figure S2

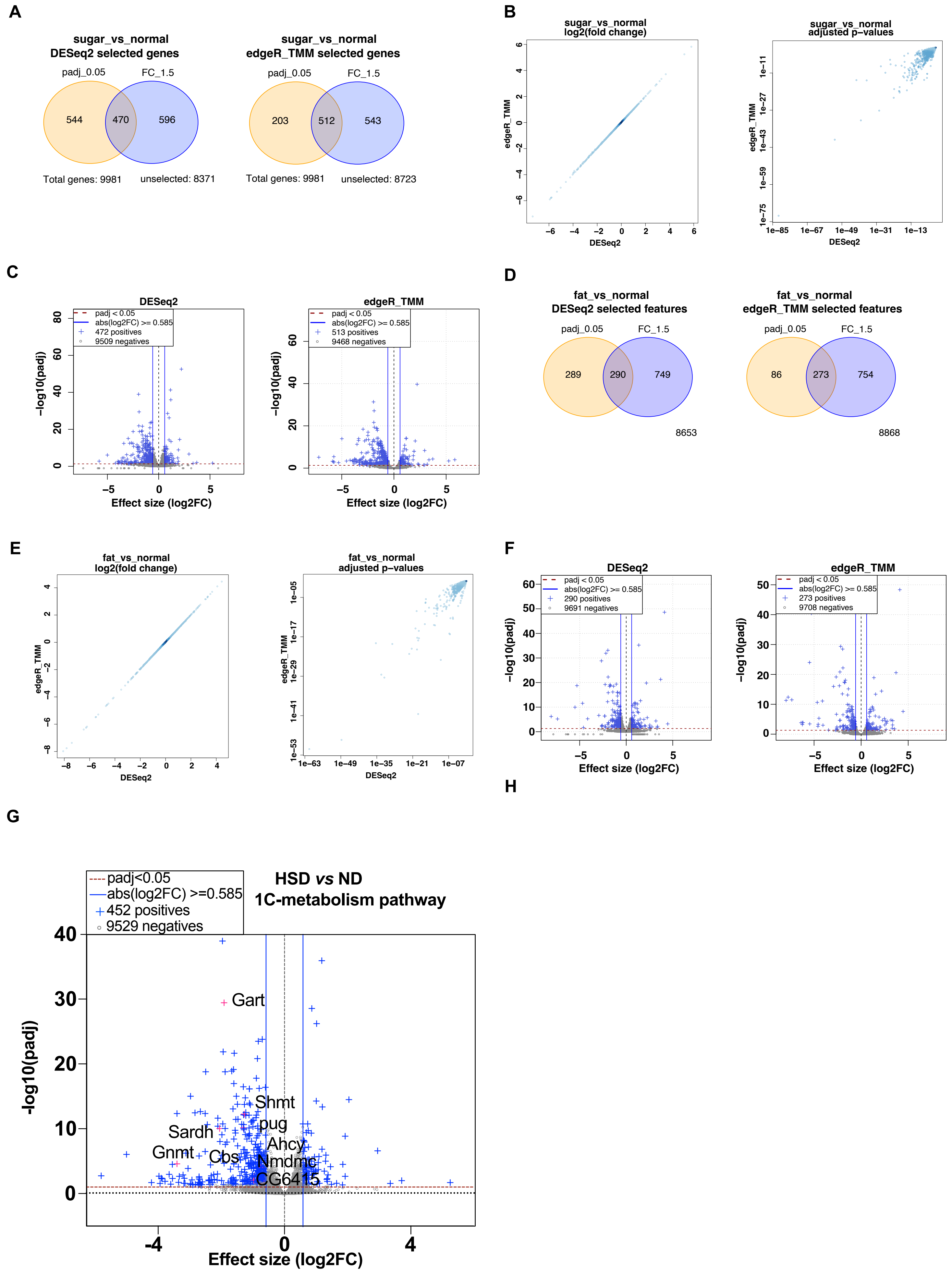

### Figure S3

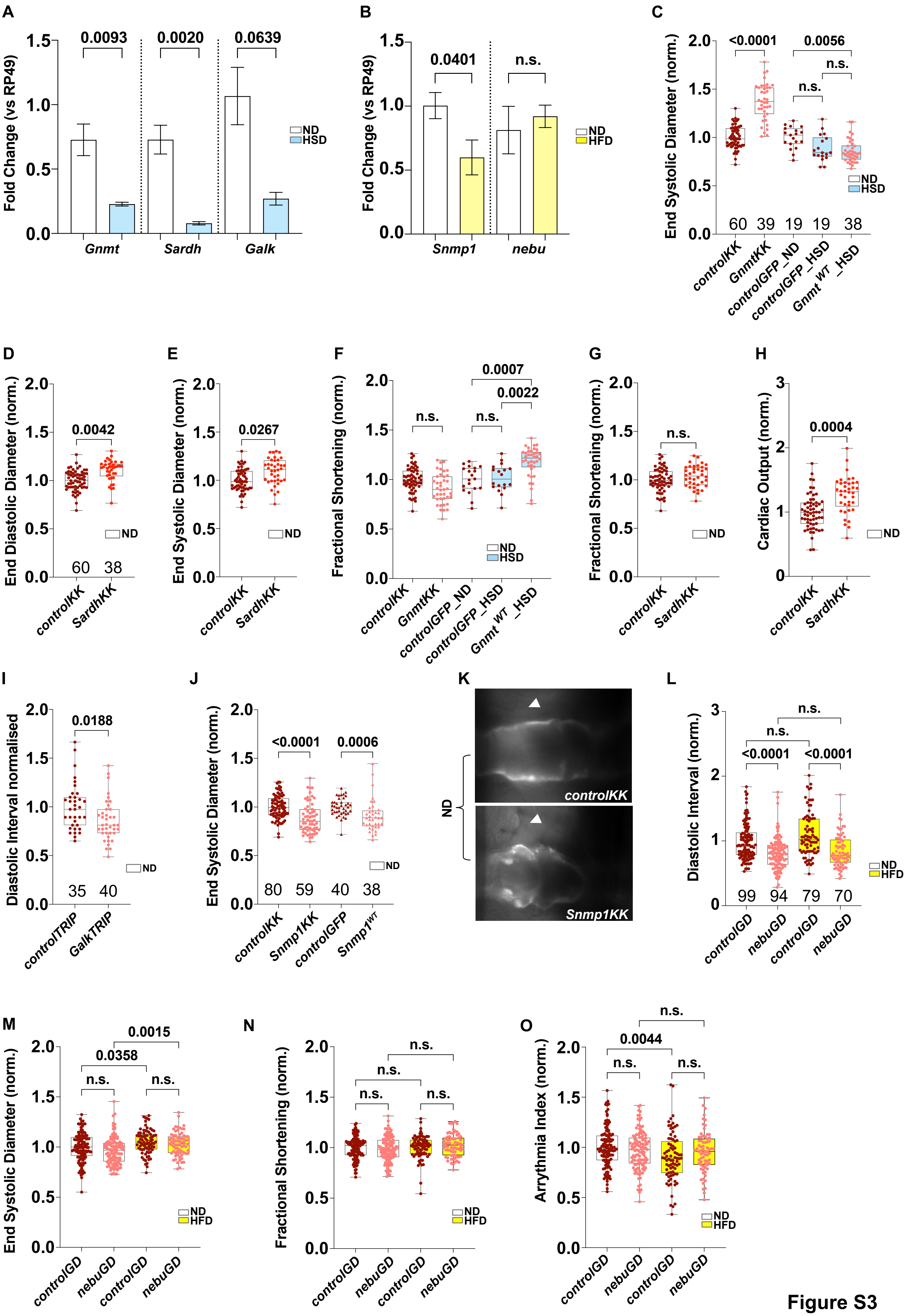

### Figure S4

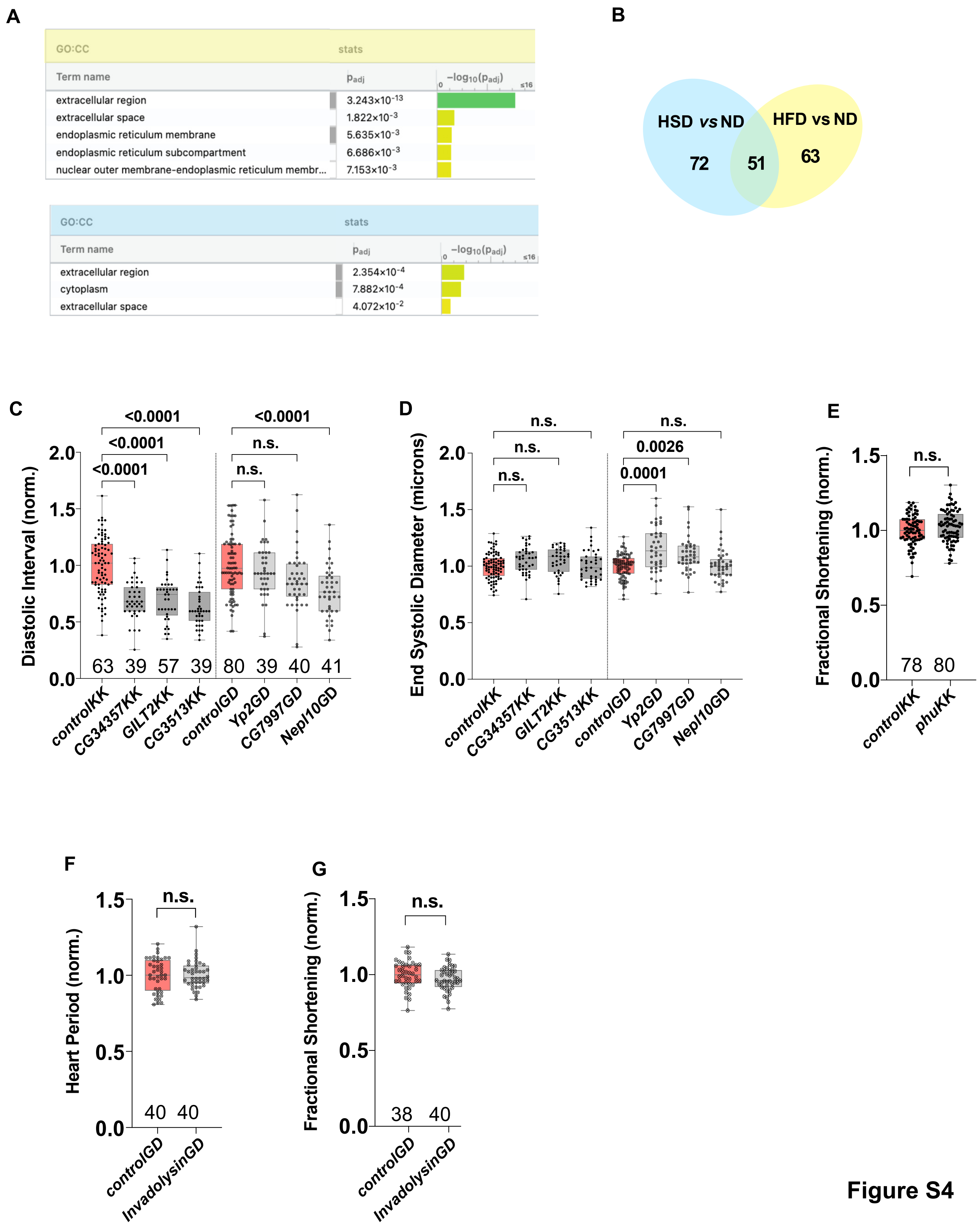

### Figure S5

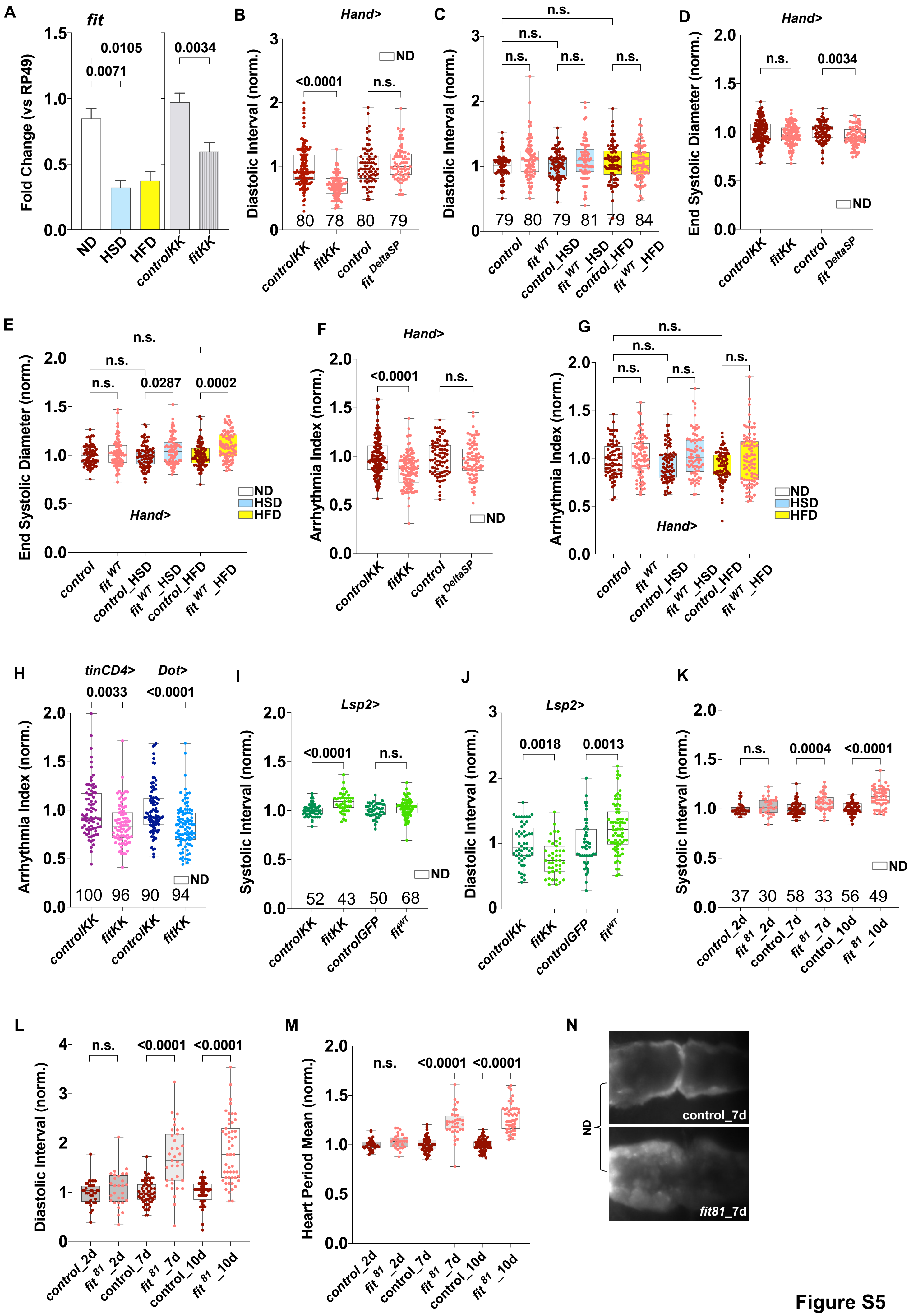

### Figure S6

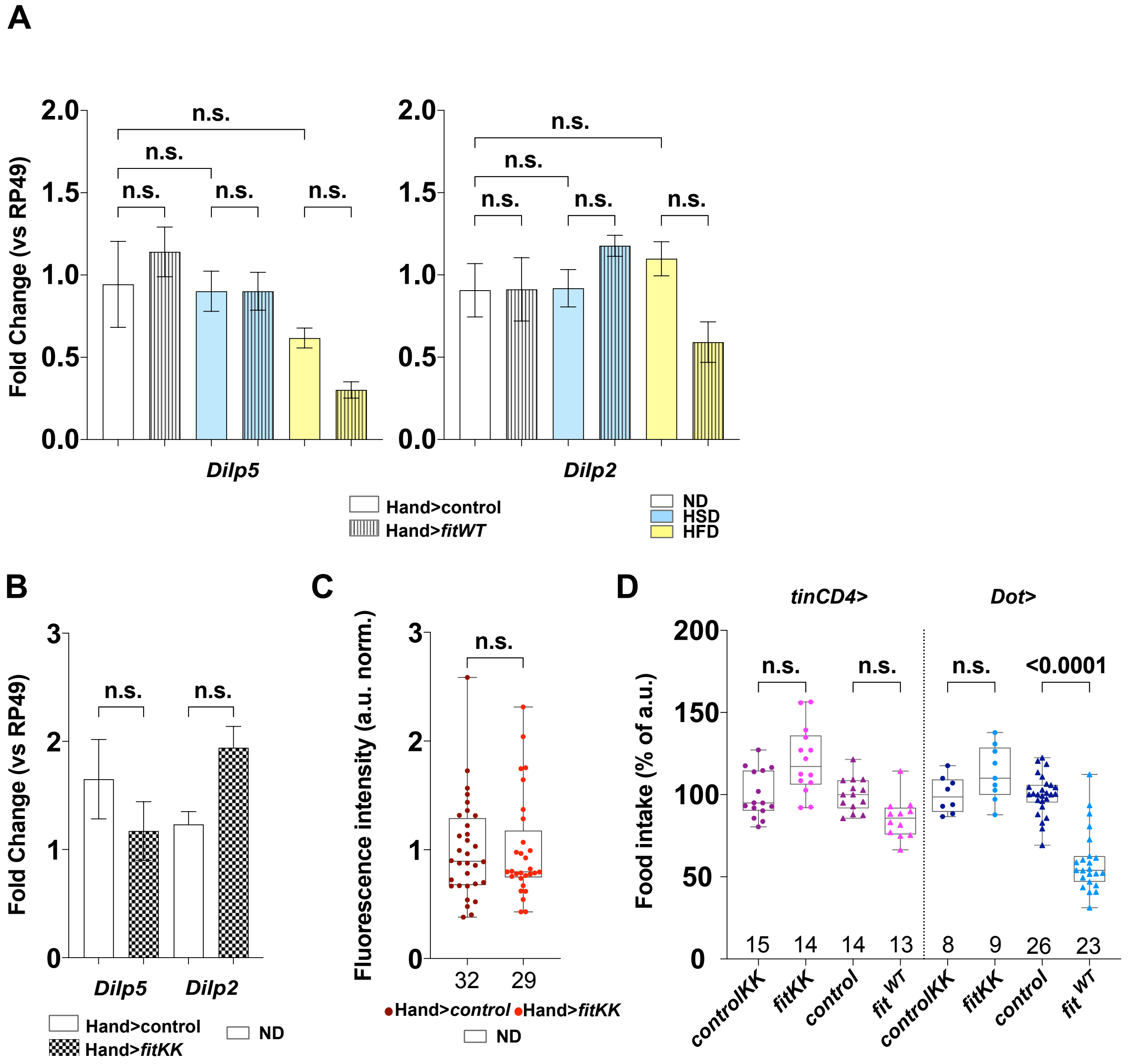
